# *De-novo* variants in *XIRP1* associated with polydactyly and polysyndactyly in Holstein cattle

**DOI:** 10.1101/574061

**Authors:** Marina Braun, Maren Hellige, Ingo Gerhauser, Malgorzata Ciurkiewicz, Annika Lehmbecker, Andreas Beineke, Wolfgang Baumgärtner, Julia Metzger, Ottmar Distl

## Abstract

Congenital polydactylous cattle are sporadically observed. Impairment of the limb patterning process due to altered control of the zone of polarizing activity (ZPA) was associated in several species with preaxial polydactyly and syndactyly. In cattle, the role of ZPA and other genes involved in limb patterning for polydactyly was not yet elucidated. Herein, we report on a preaxial type II polydactyly and a praeaxial type II+V polysyndactyly in two Holstein calves and screen whole genome sequencing data for associated variants. Using whole genome sequencing data of both affected calves did not show mutations in the candidate regions of ZRS and pZRS or in candidate genes associated with polydactyly, syndactyly and polysyndactyly in other species. Two indels, which are located in *XIRP1* within a common haplotype, were highly associated with the two phenotypes. Bioinformatic analyses retrieved an interaction between *XIRP1* and *FGFR1, CTNNB1* and *CTNND1* supporting a link between the *XIRP1* variants and embryonic limb patterning. The heterozygous haplotype was highly associated with the present polydactylous phenotypes due to dominant mode of inheritance with an incomplete penetrance in Holstein cattle.

## INTRODUCTION

Polydactyly in cattle is characterized by a variable number of supernumerary phalanges found either in all four or in single front or hind limbs. This condition can be accompanied by a complete or partial fusion of digits or by non-division of additional digits, designated as polysyndactyly (Kaus 1968). Congenital types of polydactyly have been reported in several cattle breeds including Simmentaler (Schummer 1935; Marolt and ilijas 1967), Fleckvieh (Kaus 1968), Hereford (Morrill 1945), Simmentaler-Hereford crossbreed (Johnson *et al.* 1981) and Holstein-Friesian (Marolt and Ilijas 1967; VERMUNT *et al.* 2000; Bähr *et al.* 2003; Carstanjen *et al.* 2010; Sangwan *et al.* 2015). Johnson *et al.* (1982) proposed a subdivision into seven different types (Table 1). The most common form in cattle was found to be bilateral polydactyly of both limbs with additional preaxial metacarpal bones (Johnson *et al.* 1982; MATHER 1987). In addition, polydactyly can be accompanied by further malformations like those with an additional limb (polymelia) located in different areas of the body (Kim *et al.* 2001; Murondoti *and* Busayi 2001; ALAM *et al.* 2007; FREICK *et al.* 2014). In some affected polydactylous limbs an axial rotation resulting in unequal load and lameness of different severity was seen (SchÖnfelder *et al.* 2003; Altenbrunner-martinek *et al.* 2007).

**Table 1.**
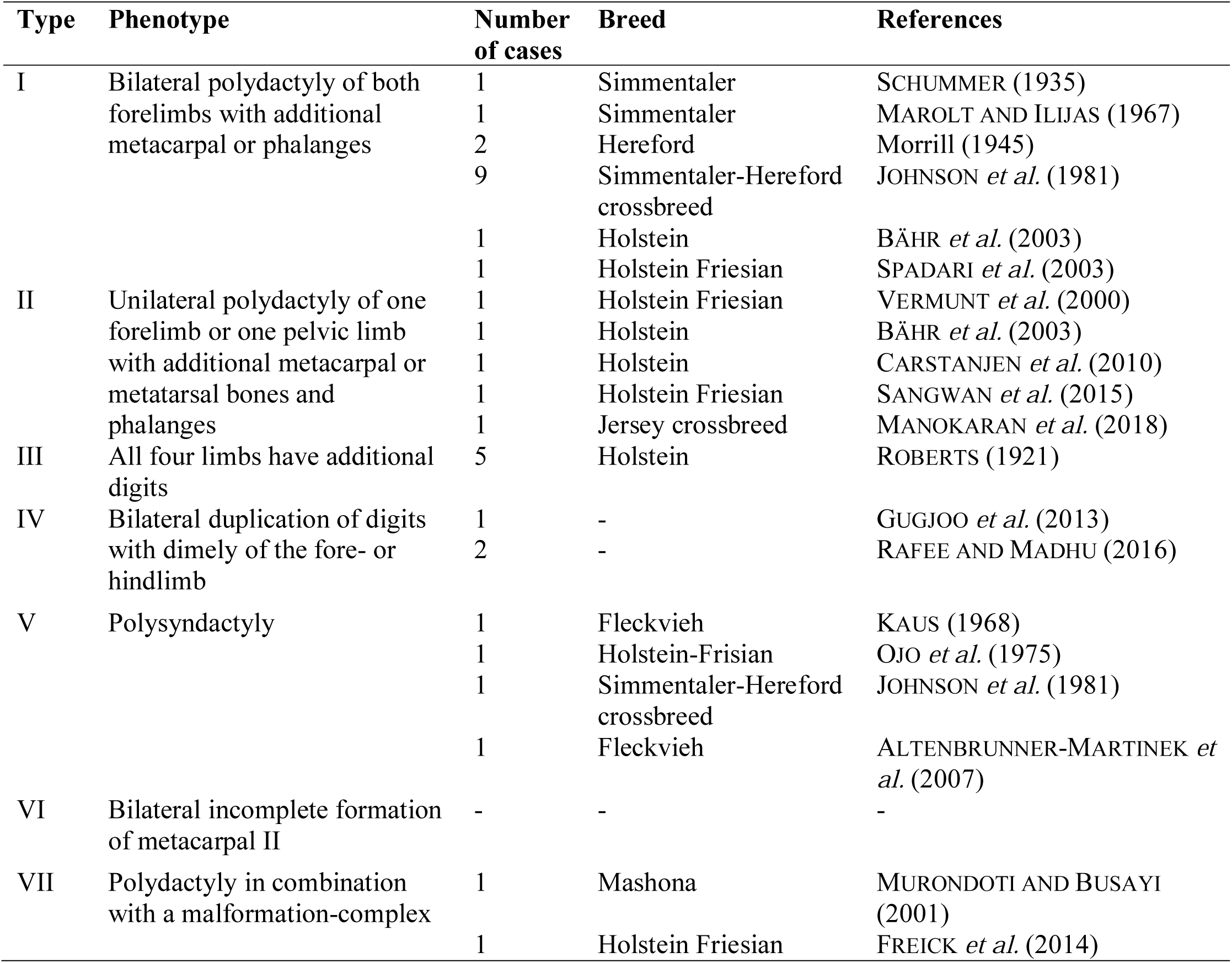
Previously reported types of polydactyly in cattle classified according to Johnson *et al.* (1982).

Different models of inheritance were suggested for bovine polydactyly like monogenic autosomal dominant (Roberts 1921) or recessive (Mather 1987), polygenic with one dominant gene linked with homozygous recessive genes on further gene loci (Johnson *et al.* 1981) and X-chromosomal recessive (Morrill 1945).

A critical candidate region for preaxial polydactyly in human and mice harbours the *sonic hedgehog (SHH)* gene (Lettice *et al.* 2003). It was postulated that supernumerary digits develop during limb patterning along the anterior–posterior (AP) axis, controlled by a cell region of the limb bud designated as the zone of polarizing activity (ZPA). This region was shown to express at high amounts the *SHH* gene involved in embryonic bone development and remodeling, specification of the digit number and the digit identity (Ingham AND Mcmahon 2001). In addition, further control of ZPA and *SHH* is achieved by a long-range cis-regulatory element, designated as the ZPA regulatory sequence (ZRS) which is located at about 1 megabase upstream from *SHH* within intron 5 of the *limb development membrane protein 1* gene (*LMBR1*) (Laurell *et al.* 2012). Studies in mice and chicken affected with preaxial polydactyly verified altered regulatory functions of ZRS (Masuya *et al.* 2007; Maas *et al.* 2011). More than 20 different mutations in the ZRS and in the upstream sequence of the ZRS (pZRS) have been identified to be associated with polydactyly in human (Lettice *et al.* 2003; Gurnett *et al.* 2007; Furniss *et al.* 2008; Semerci *et al.* 2009; Wieczorek *et al.* 2009; Farooq *et al.* 2010; Albuisson *et al.* 2011; AL-QATTAN *et al.* 2012; Laurell *et al.* 2012; Vandermeer *et al.* 2012; Girisha *et al.* 2014; Norbnop *et al.* 2014; Vandermeer *et al.* 2014; Wu *et al.* 2016; Xiang *et al.* 2017; Hovius 2018), dogs (Park *et al.* 2008), cats (Lettice *et al.* 2008) and chicken (Huang *et al.* 2006; Dorshorst *et al.* 2010; Dunn *et al.* 2011; Johnson *et al.* 2014) so far. In human, limb malformations caused by mutations located in this regulatory region represent a wide range of phenotypes (Vandermeer and Ahituv 2011). In cattle, the role of *SHH*, ZRS and pZRS for polydactyly has not yet been investigated for variants associated with a polydactylous condition.

Therefore, the objective of this study was to characterize the phenotypes of two polydactylous Holstein calves and screen whole genome sequencing data for all genomic candidate regions in mammalian species and all private variants for these calves. In addition, cytogenetic analysis for chromosomal aberrations and search for structural genomic variants were performed.

## METHODS

### Animals

Two polydactylous Holstein calves born on different dairy farms in Germany were examined for this study. Complications or health problems during gestation and calving difficulties were not reported for the dams. Both cases were transferred to the Institute for Animal Breeding and Genetics of the University of Veterinary Medicine Hannover for further examination. The first calf (case 1) was born without complications in September 2016 after a normal gestation period. According to the report of the farmer, the parents of the calf had no signs of visible malformations. The calf was subsequently euthanized due to humanity reasons because of severe lameness and its inability to stand. Euthanasia was performed by intravenous injection of ketamine combined with xylazine as premedication following by pentobarbital-natrium (Release®). Case 2 was a male Holstein calf born in March 2018. The dam and the sire of the calf had no visible abnormalities. The dam had given birth to a normal calf two years before. This maternal half-sister of the calf born two years earlier showed no polydactyly or any other abnormalities. The affected calf was euthanized by the veterinary practitioner and transferred for further examinations to the Institute for Animal Breeding and Genetics of the University of Veterinary Medicine Hannover. We collected EDTA-blood samples from the two affected calves, their dams and one maternal half-sibling of case 2 as well as sperm samples from the sires.

All animal work has been conducted according to the national and international guidelines for animal welfare. Sampling from the cases and healthy controls were approved by the Institutional Animal Care and Use Committee of Lower Saxony, the State Veterinary Office Niedersächsisches Landesamt für Verbraucherschutz und Lebensmittelsicherheit, Oldenburg, Germany (registration number 33.9-42502-05-04A247). Both animals were euthanized because of the bad health condition resulting from the unability to stand and drink without support and the steadily declining accompanying risk of infections. The decision to euthanize the animals was at the discretion of the supporting veterinarians. The cattle owners agreed that the animals and their samples are part of our study and a written informed consent was given for publication of this research article and any accompanying images.

### Clinical and radiological examination

Both affected calves underwent a clinical examination. For case 1, computed tomography (CT) was performed post mortem in sternal recumbency position. CT scans were acquired with a multislice helical CT scanner (Brilliance TM CT 16 BigBore, Philips Healthcare, Netherlands) using conventional settings (120 kV and 315 mA) and 0.8 mm (0.031 inch) slice thickness.

### Necropsy and patho-histological examination

The two affected calves underwent necropsy after euthanization. Malformed forelimbs were macerated to characterize the osseous malformations in detail. Samples of the bones (femur and sternum), muscles (diaphragm, *M. quadriceps femoris*) and all organs were fixed with 10% formalin, and embedded in paraffin. Sections of 4 μm were stained with eosin and hematoxylin (HE) and histologically screened for morphological changes.

### Pedigree analysis

The pedigree data of both affected calves were used to calculate the relationship and inbreeding coefficients based on the maximal depth of eleven generations with OPTI-MATE version 4.0 (Schmidt *et al.* 2006). In addition, contributions of ancestors to the inbreeding coefficients were determined.

### Cytogenetic analysis

Cytogenetic analysis were performed to identify chromosomal aberrations in the two polydactylous calves. The chromosome preparation followed standard protocols using heparin blood samples for lymphocyte cultures preparing (Wockl *et al.* 1980). The metaphase chromosome were stained with giemsa and visualized using the microscope Axioplan 2 (Carl Zeiss Microscopy, Jena, Germany). Photos were taken with a computer-controlled CCD camera, Axiocam 105 color (Carl Zeiss Microscopy), and processed with ZEN 2 (blue edition) software (Carl Zeiss Microscopy). In total, 50 metaphase chromosomes were analyzed.

### Whole genome sequencing

For WGS, genomic DNA of both affected calves and the dam of case 2 was isolated using standard chlorophorm extraction. After preparing libraries from these samples according to the manufacturer’s protocols using the NEBNext Ultra DNA Library Prep Kit for Illumina (New England BioLabs, Ipswich, MA, USA), WGS was performed using a NextSeq500 (Illumina, San Diego, CA, USA) in a 2 x150 bp paired-end mode. Data were controlled for quality using fastqc 0.11.5 (Andrews 2015) and the reads were trimmed using PRINSEQ, version 0.20.4 (Schmieder and Edwards 2011). Data were mapped to the bovine reference genome UMD 3.1. (Ensembl) using BWA 0.7.13 (Li and Durbin 2009). Sorting, indexing and marking of duplicates of Bam-files were performed using SAMtools 1.3.1 (LI *et al.* 2009) and Picard tools (http://broadinstitute.github.io/picard/, version 2.9.4). Variants were called with GATK, version 4.0 (Mckenna *et al.* 2010), using Base Quality Score Recalibration (BQSR), Haplotype Caller and Variant Recalibrator. All variants with a read depth of 2-999 and quality score values >20 were chosen from the vcf-file for further filtering. Variants of the two polydactylous calves and the dam of case 2 were compared with the vcf files of 90 private controls including cattle from the breeds Holstein, Fleckvieh, Braunvieh, Vorderwald, German Angus, Galloway, Limousin, Charolais, Hereford, Tyrolean Grey and Miniature Zebu.

Only those variants with high or moderate effects according to prediction toolbox SNPEff, version 4.3 q (2017-08-30, SNPEff database UMD3.1.86) (Cingolani *et al.* 2012) were selected. Homozygous or heterozygous mutant variants in each of the two affected Holstein calves and heterozygous mutant or homozygous wild type in the dam but homozygous wild type in all 90 control individuals were filtered out using SAS, version 9.4 (Statistical Analysis System, Cary, NC, USA). These variants were investigated for their potential functional effects and compared with candidate genes for polydactyly, syndactyly and polysyndactyly known in human and domestic animals (Table S1) according to National Center for Biotechnology Information (NCBI, www.ncbi.nlm.nih.gov) and Online Mendelian Inheritance in Animals (OMIA, www.omia.org, date of access). To verify the previously investigated protein effect of the remaining variant, we applied the *PolyPhen-2* (Polymorphism Phenotyping v2) tool. The position of the variants in the UMD3.1 genome were compared with the genome database ARS-UCD1.2 using the ncbi genome remapping service (NCBI, www.ncbi.nlm.nih.gov/genome/tools/remap). Variants with discording genomic position were omitted and variants which had the same genotype in the affected calves and in the dam were also excluded, excepted for one variant which was located in a common gene next to another critical variant.

To define the bovine ZRS and pZRS regions, multiple sequence alignments were performed for the bovine region (UMD3.1 and ARS-UCD1.2) of intron 5 of *LMBR1* with ZRS and pZRS regions in human, mouse, chicken, dog and cat using ClustalW2, version 2.1 (Larkin *et al.* 2007). Previously reported mutations associated with polydactyly of each sequence were flagged. We screened the WGS data of both affected calves for mutant variants in this matched region of intron 5 using SAS, version 9.4.

In the next step, structural variants were retrieved with GASVPro (Geometric Analysis of Structural Variants Pro), version 1.2.1 (Sindi *et al.* 2012) and LUMPY (LAYER *et al.* 2014) and compared in-between the affected calves. We analyzed bam-files derived from WGS data of the affected calves, the dam of case 2 as well as 12 controls to screen for structural variants potentially involved in the development of polydactyly. A scan for all possible breakpoints or breakends by integrating read depth and paired read signals into a unified probabilistic model was employed (Sindi *et al.* 2012). The final list of predictions was pruned for redundant predictions and filtered for significant events. The remaining structural variants were compared in-between the affected calves and the controls separately for GASVPro and LUMPY as well as cross-validated among GASVPro and the breakpoint prediction framework LUMPY.

### Validation

The identified candidate variants g.118035410_128del within the gene *ceramide kinase* (*CERK*) and g.12552417dupC and g.12552582delA) within the gene *xin actin binding repeat containing 1* (*XIRP1*) were validated in the two affected calves, their parents and the maternal half-sibling of case 2 using PCR amplicon and Sanger sequencing. Genomic DNA was extracted using standard ethanol fraction. We designed primer pairs with Primer3 tool, version 0.4.0 (www.bioinfo.ut.ee/primer3-0.4.0/), for PCR amplicon (Table S2). To visualize predicted changes in the *XIRP1* protein, protein sequences of the wild type protein (ENSBTAP00000054446) and predicted mutant proteins were compared and aligned using ClustalW2, version 2.1. Genetic interactions between *XIRP1* and critical candidate genes for the polydactylous phenotype were investigated using GeneMANIA (Warde-farley *et al.* 2010).

For genotyping 359 unrelated animals with an unknown phenotype of the breeds Holstein (n=276), German Fleckvieh (n=12), German Brown (n=12), Charolais (n=12), Limousin (n=11), Blonde d’Aquitaine (n=12) and Salers (n=12), we designed a restriction fragment length polymorphisms (RFLP) for each variant (Table S2). For the deletion (g.12552582delA), we used the same primer pairs as for validation and for the duplication (g.12552417dupC) we created a mismatch primer.

For WGS data validation of the ZRS and pZRS regions by Sanger sequencing, three primer pairs were designed covering both regions (Table S3) using Primer3 tool, version 0.4.0.

### Availability of data and materials

WGS data of the two calves (case 1 and 2) and the dam of case 2 one were deposited in NCBI Sequence Read Archive (Submission ID: SUB4556003, BioProject PRJNA492844 with IDs *SAMN10108008, SAMN10108011, SAMN10108012)*.

## RESULTS

### Clinical examination

The calves showed a unilateral polydactyly of the right forelimb (Figure 1AB, 2AB). The polydactylous forelimb of case 1 had an additional remarkable angulation to the medial side. This calf had problems to stand for a longer period. In the beginning of standing, the calf was able to stretch the limb for a while. Then the limb bent in the carpal joint. While walking it showed a severe lameness and was unable to put its full weight on the right front limb. Therefore, it limped by making a short step and fell subsequently back to the non-affected left limb. Case 2 had no problems to stand and walk and no significant lameness was detected. The polydactylous claws had one additional digit in each calf, but the arrangement was different among calves. The affected claws of case 1 were in an abnormal angled position. They were medially angled. The medial one had a straight direction and the claw surface could move on the ground. On contrast, the lateral claw was flexed backwards. Distally to the metacarpal region there was one dewclaw dorsally to the lateral claw. The structure of the claws was physiological. The additional claw was located at the palmar aspect of the angled metacarpal region. It showed a separate claw which was not fused with one of the others. The slightly blunt cone-shaped form was rugged at its corneal layer. The additional digit of case 2 was medially located and partially fused together with the other digit next to. Due to the fusion, all three claws formed a common enlarged sole without rotation. Case 2 showed also a cleft palate which caused nasal reflux accompanied by cough after suckling milk. Further, on the dorsal aspect of the carpal joint of case 1, a circular defect of the skin was notified.

**Figure 1.**
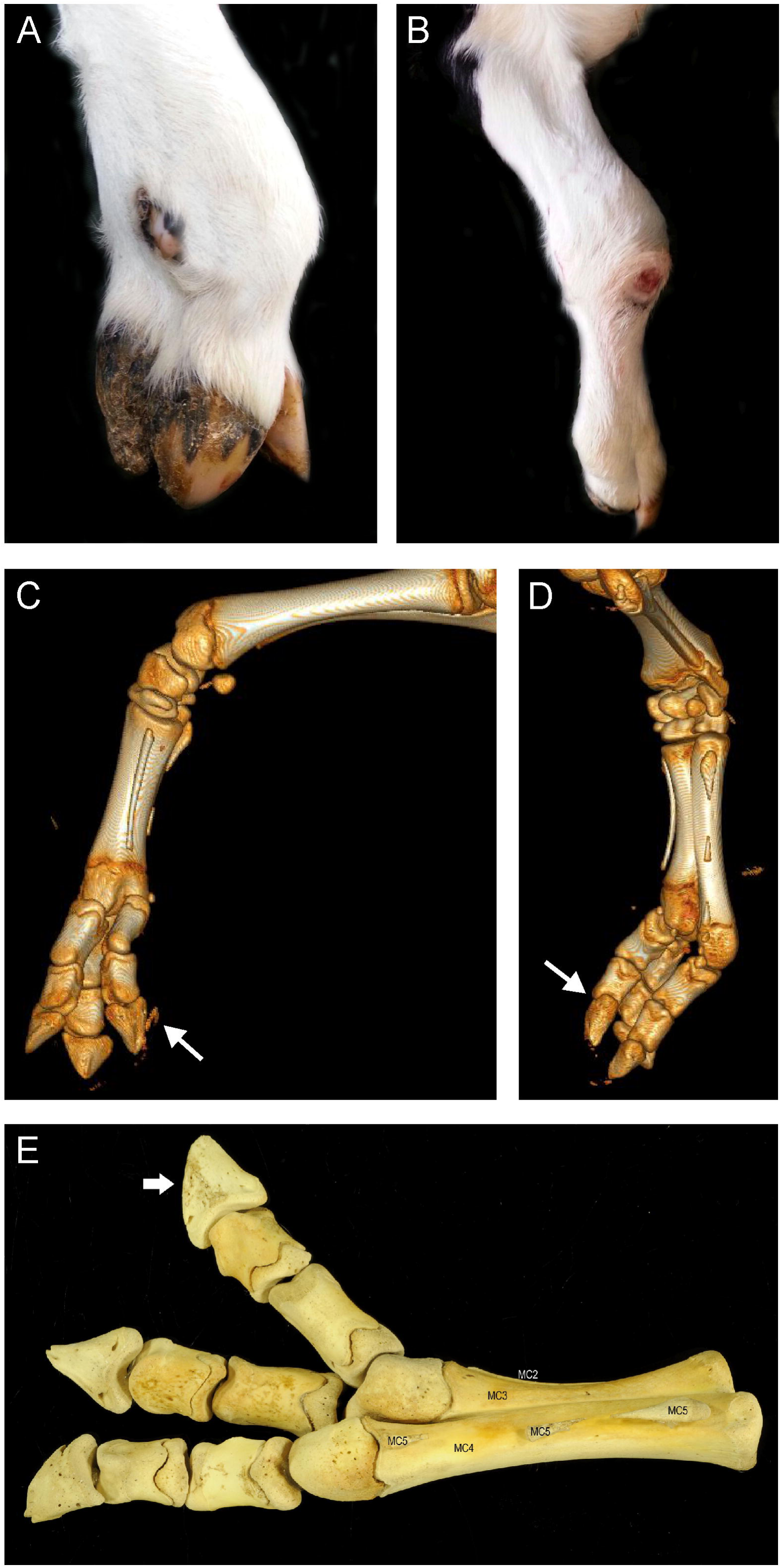
The polydactylous right forelimb of case 1 in the **a** lateral and **b** dorsal view. Computed tomography (CT) image acquisition of the right front limb in **c** lateral and **d** palmar view. The additional digit is marked with an arrow. **e** Lateropalmar view on the macerated right forelimb shows the formation of an additional digit (arrow) with three phalanges, an additional articular surface located at palmar side of the third metacarpal bone (MC3) and the formation of a fifth metacarpal bone (MC5) composed of three separate segments as well as a second metacarpal bone (MC2).

### Computed tomographic examination

The lateral half of the distal epiphysis of the radius was shortened (Figure 1CD). The third and fourth metacarpal bone were fused in the proximal half of the right forelimb and the incomplete bony septum in-between was thickened compared to the contralateral thoracic limb. An additional metacarpal bone II was present with a length of 8 cm (3.15 inches). The distal heads of the metacarpal bones were misshaped. The lateral metacarpophalangeal joint was angulated and the fourth metacarpal bone was elongated in comparison to the third. A proximal, middle and distal phalanx of the digit IV were present and the proximal sesamoid bones were displaced. The third digit was parallel to the fourth and placed at the medial border of the head of the third metacarpal bone. There was an additional digit with proximal, middle and distal phalanx in contact with the third metacarpal bone. This additional digit had an articulation with the misshaped head of the third metacarpal bone at the palmar aspect.

### Necropsy and patho-histological examination

The right forelimb of case 1 demonstrated a formation of an additional digit with three phalanges and associated proximal and distal sesamoid bones, which articulated with an additional articular surface located at the palmar side of the third metacarpal bone (MC3). The fused third and fourth metacarpal bones differed in length resulting in a 1.8 cm (0.71 inch) protuberance of the distal epiphysis and a 0.5 cm (0.19 inch) protuberance of the proximal epiphysis of MC4. Both normal digits were rotated by 30° laterally. In addition, a fifth metacarpal bone (MC5) composed of three separate segments and a second metacarpal bone (MC2) were present (Figure 1E). The tendon of the *M. flexor digitalis profundus* inserted at the lateral normal and the additional digits instead of the two normal digits. Similarly, the tendon of the *M. interosseus* inserted at the proximal sesamoid bones of the lateral normal and the additional digits instead of the proximal sesamoid bones of the two normal digits.

The right forelimb of case 2 showed medially a polysyndactyly with three digits showing a partial fusion (Figure 2AB). All digits had regularly formed and completely separated distal and medial phalanges, while the proximal phalanx of the medial digits was partially fused (Figure 2D-G). The fusion extended over the entire metaphysis and the palmar aspect of the epiphyses. On the dorsal aspect, the proximal and distal epiphyses were partly separated by longitudinal grooves. Distally, two articular surfaces were present for the medial phalanges. Proximally, two shallow joint sockets were present that articulated with the medial metacarpal bone. The metacarpus consisted of two asymmetrical bones, fused at the center of the metaphyses. The medial metacarpal bone was broader and longer than the lateral one, resulting in a 0.6 cm (0.24 inch) protuberance of the distal epiphysis and a 0.3 cm (0.21 inch) protuberance of the proximal epiphysis. The epiphyses and the distal and proximal ends of the metaphyses lacked fusion. An additional, rudimentary metacarpal bone was present at the caudal aspect of the metacarpus, showing a fusion with the lateral metacarpal bone.

**Figure 2.**
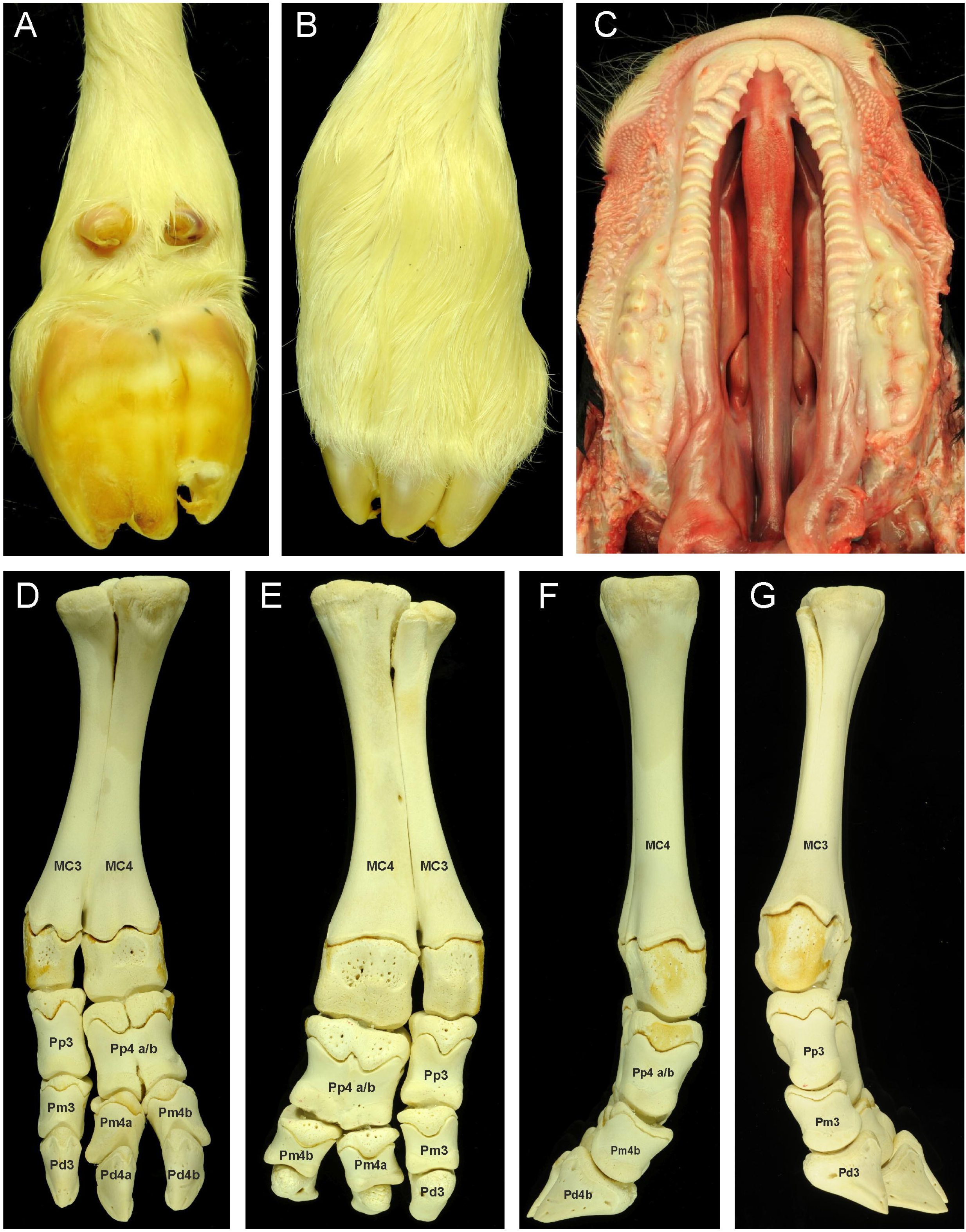
The poly-and syndactylous right forelimb of case 2 in **a** palmar and **b** dorsal view. **c** The prominent cleft palate 2 including the hard and soft palate. **d** Dorsal, **e** palmar, **f** medial and **g** lateral view on the macerated right forelimb shows three digits with separated distal and medial phalanges and a fuses proximal phalanx of the medial digits. An additional metacarpal bone is present on the palmar.

In addition to the malformations of the forelimb, case 2 exhibited a cleft soft and hard palate in the oral cavity (Figure 2C). The medial fissure began behind the incisors and was extended for 17 cm (6.69 inches) in length and 3 cm (1.18 inch) wide, to the pharynx.

Histopathological changes were not detected in any of the examined organs.

### Pedigree analysis

Case 1 had a pedigree completeness of 90.3% and 45.8% over 5 generations and 11 generations. Pedigree completeness for case 2 was 96.7% over 4 generations and 34.27% over 11 generations. The relationship coefficient between the two affected calves was 4.95%. Inbreeding coefficients for case 1 and case 2 were 2.91 and 0.71%, respectively. The bulls Chief (18.79%), Valiant (15.0%) and Elevation (14.5%) mainly contributed to the inbreeding coefficients for both calves.

### Cytogenetic analysis

The microscopic presentation of the metaphase chromosomes displayed in both calves a normal karyogramme with 2n= 60 chromosomes. Numerical aberrations of autosomal and sex chromosomes in the female and male calf were not observed in any of the metaphases screened or chromosomal aberrations.

### Whole genome sequencing

Filtering WGS data (Table 2) displayed 259 variants in case 1 and 248 variants in case 2. No protein coding variant was located in critical candidate genes. Four heterozygous variants were identified, which are exclusively shared by both affected calves and two further heterozygous variants shared by the two cases and the dam of case 2 (Table S4). Among these six variants, we identified three critical candidate variants *CERK*:g.118035410_128del, *XIRP1*:g.12552417dupC and *XIRP1*:g.12552582delA.

**Table 2.** Filtering process of whole genome sequencing variants for both affected calves. The resulting variants after each step are given for case 1 case 2.

| Filtering step | Resulting variants |  |
| --- | --- | --- |
|  | Case 1 | Case 2 |
| Variants with a read depth of 2-999 and quality score values >20 | 37,707,104 | 37,760,993 |
| Comparing variants with high or moderate effects to 90 private control animal | 259 | 248 |
| Comparing variants with candidate genes in other species | 0 | 0 |
| Comparing variants in-between the affected calves | 6 | 6 |
| Comparing UMD3.1 variant position with ARS_UCD1.2 and omitting variants with discording genomic position | 4 | 4 |
| Excluding variants showing the same genotype as the dam | 2 | 2 |
| Retake one variant locating in one of the final genes | 3 | 3 |
| Validation and excluding variants occurring in the same genotype in the unaffected and affected animals | 2 | 2 |

Examination of filtered WGS data of each polydactylous calf revealed no polydactylous-associated variants in the bovine ZRS (Figure S1) and pZRS (Figure S2) regions. These candidate regions are located within intron 5 of *LMBR1* (ENSBTAG00000030817) (Figure S3). The bovine sequence of the ZRS and pZRS showed a high percentage of base pairs matching.

The results from data analysis using structural variant detecting software did not reveal structural variants exclusively found in both polydactylous calves.

### Validation

The *CERK* variant occurred heterozygous in both affected calves, but also in the unaffected maternal half-sibling and sire of case 2. This excluded this variant for further examination and genotyping. Validation of the two candidate variants in *XIRP1* (Figure 3) through Sanger sequencing of familiar members revealed both calves as heterozygous mutant for both variants (Figure 4) as well as the dam of case 2. Both sires (sperm DNA) and a maternal half-sibling of case 2 were homozygous wild type for both variants (Figure S4). The *XIRP1* variant p.Ala1076fs was predicted to create a frameshift and thus a new protein sequence starting with a prolin instead of an alanine (Figure 5). The protein sequence gets back in frame by the G duplication (p.Trp1130fs) after 54 amino acids (p.1076 – p.1130). Both frameshift variants were predicted to be probably damaging (p.Ala1076fs, 0.998) and possibly damaging (p.Trp1130fs, 0.922).

**Figure 3.**
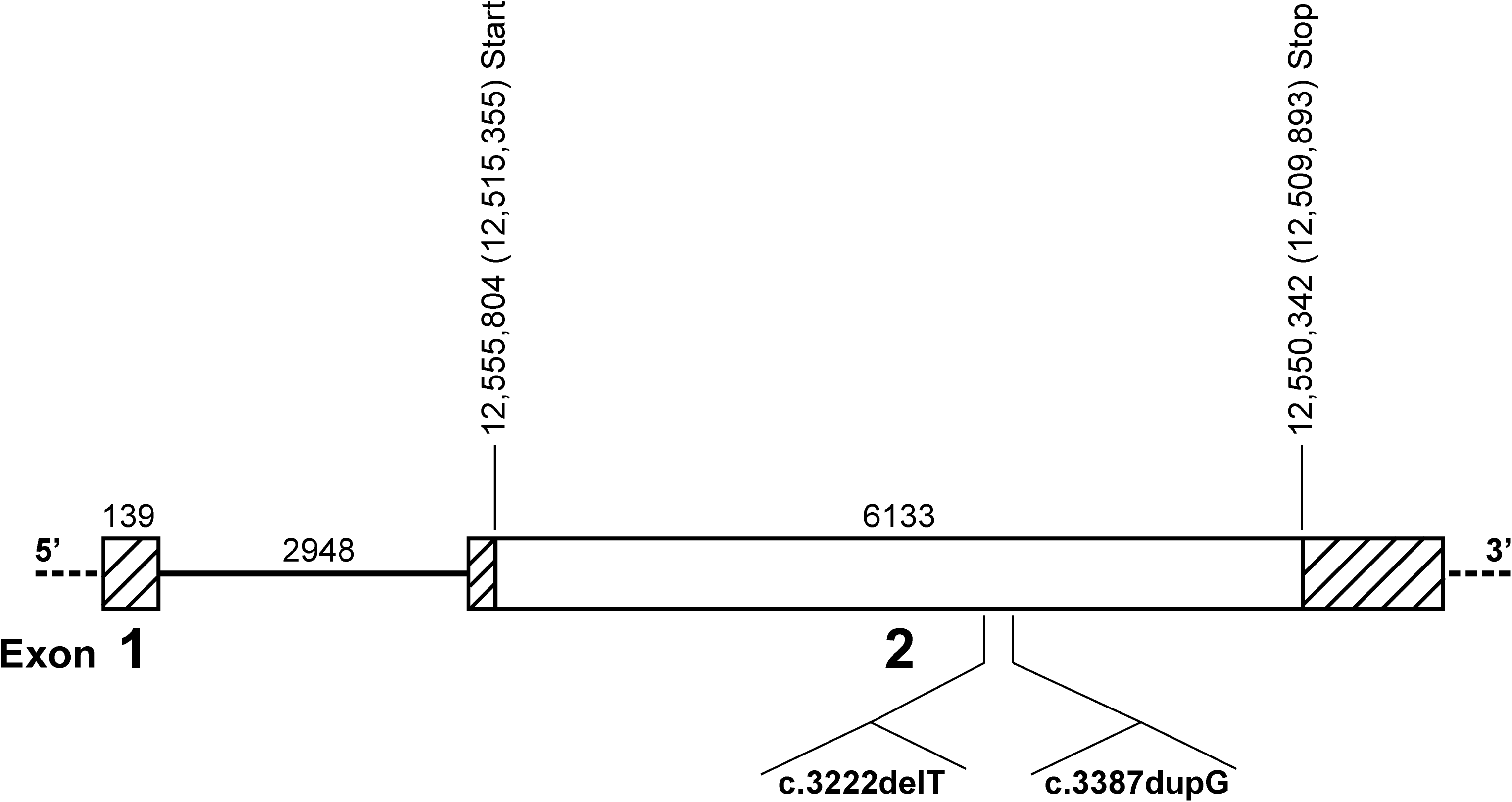
Gene model of the bovine *XIRP1* (ENSBTAG00000046512) gene. The location of the filtered variants (g.12552417dupC, g.12552582delA) in the coding region of exon 1 are marked.

**Figure 4.**
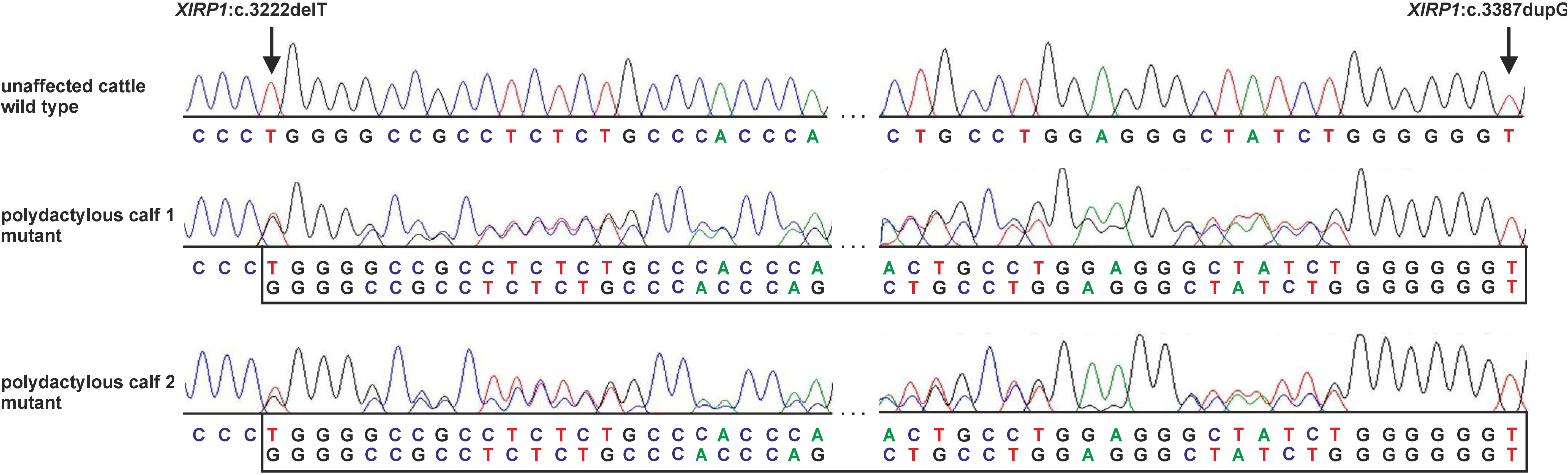
Sanger sequence of the heterozygous occurrence of both filtered variants in *XIRP1* in the two affected calves compared to the sequence of one control animal.

**Figure 5.**
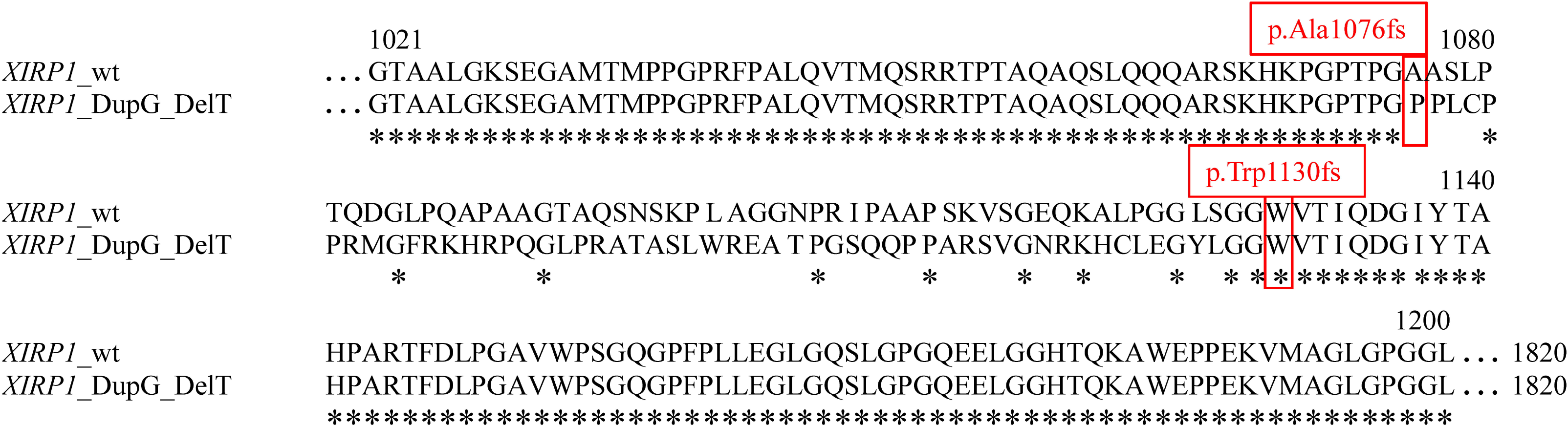
Variant effects on *XIRP1* protein. The wild type (wt) amino acid sequence of cattle transcript *XIRP1*-201 (ENSBTAP00000054446) is displayed. The modifications of the amino acid sequence caused by the combination of duplication (c.3387dupG) and the deletion (c.3222delT) predicted a change of 54 amino acids between p.1076 and p.1130. Identical amino acids between the protein sequences are marked with asterisks.

A direct genetic interaction between *XIRP1, FGFR1, catenin beta 1* (*CTNNB1*) and *catenin delta 1* (*CTNND1*) as well as indirect interactions in-between *LMBR1 Domain-Containing Protein 1* (*LMBRD1*) and *SHH* and *LMBR1* were predicted using GeneMANIA.

Genotyping of 359 animals revealed in Holstein cattle eight heterozygous and one homozygous carriers of the two variants g.12552417dupC and g.12552582delA (Table S5). None of the two variants was detected in the 83 individuals of the other six cattle breeds.

Sanger sequencing of the candidate region in intron 5 of *LMBR1* orthologues to human, mouse, chicken, dog and cat ZRS and pZRS confirmed the assumption of absent variants.

## Discussion

In this study, we identified two *de-novo XIRP1* variants associated with polydactyly and polysyndactyly in Holstein cattle. The investigated phenotypes were assigned to a preaxial type II polydactyly and praeaxial type II+V polysyndactyly. These types have already been suggested to a hereditary condition in polydactyly types I to VI (Johnson *et al.* 1982). Various classifications found in previous reports supported this assumption, proposing the unilateral polydactyly type II to be the most common form in the Holstein breed (Vermunt *et al.* 2000; BÄHR *et al.* 2003; Carstanjen *et al.* 2010; Sangwan *et al.* 2015). We found evidence that the affected calves had common ancestors, Valiant and Chief, which are known to have a high impact on Holstein bloodlines (Krugljak 2014).

Both calves had also in common that the right forelimb exhibited one additional digit. This may suggest a skeletal form of atavism, represented by additional bones found in earlier evolutionary specimens, reappearing in these calves (Tyson *et al.* 2004).

Genetic variants causative for such type of polydactyly were frequently found in the ZRS and pZRS regions located in intron 5 of *LMBR1, FGFR1* as well as *SHH* in human (Wieczorek *et al.* 2009; Xiang *et al.* 2017; Hovius 2018) and mice (HAJIHOSSEINI *et al.* 2004). In our analysis, we confirmed the high conservation of ZRS and pZRS in cattle, but excluded polydactyly-associated variants in these regions and other polydactyly-related candidate genes reported for species other than cattle.

Our WGS analysis let us suggest, that two *de-novo* variants within a haplotype of *XIRP1* are associated with polydactyly in both investigated cases. This haplotype was absent in the dam and sire of case 1 and thus these two variants arose during the zygotic stadium or in the maternal germline. In the second case, we suggest a late post-zygotic development of the same two de novo variants in the embryonic development of the dam to be most probable as it has also been found for an important fraction of de novo mutations in human (Acuna-hidalgo *et al.* 2015). This leads to the assumption that a somatic mosaicism resulted in an apparently unaffected dam but was transmitted into the calf triggering the development of additional digits as it was described in patients by Sicca *et al.* (2003) and Mineyko *et al.* (2010). Such parental somatic mosaicism has been reported to be an often underrecognized cause for the recurrence risk of genomic disorders in human (Campbell *et al.* 2014). Nevertheless, the sires of the affected calves were excluded as carriers for these variants in the germ cells, according to our validation analysis in DNA of their sperm cells and due to their widely dispersed use for insemination in Holstein. We assume, that somatic mosaicism have been occurred in the Holstein population and have been transmitted to offspring, as we were able to identify nine individuals with unknown phenotype in the validation sample, harboring the mutant *XIRP1* haplotype. It leads to the suggestion that the present phenotypes were caused by a dominant mode of inheritance of the two variants with an incomplete penetrance of the phenotype. The low frequency of mutant alleles in the Holstein population and missing mutant alleles in other breeds are supportive for a de-novo somatic mosaicism.

The two identified *de-novo* variants were predicted to be both frameshift variants presumably resulting in a shortened XIRP1 protein and thus a nonsense mediated Mrna decay, as it is usually performed for false functional or non-functional shortened proteins (Credille *et al.* 2005). Nevertheless, due to the haplotype-compared heterozygous genotype in the affected calves, we propose that the resulting protein is not shortened but modified for 54 amino acids, because the second more distal variant brings the reading frame back into the original frame and possibly prevents the modified XIRP1 mRNA from decay.

The gene *XIRP1* is important for embryonic development of skeletal muscles and heart (Grosskurth *et al.* 2008; Feng *et al.* 2013). Studies of bovine granulosa cells suggested a correlation between *XIRP1* and *fibroblast growth factor* (*FGF*) genes (Guerrero netro 2014). FGF signaling plays an important role in embryonic development and in skeletal formation (Thisse and Thisse 2005). In polydactylous chicken, interactions between the ZRS region, *SHH* and the FGF pathway signaling were confirmed (Johnson *et al.* 2014). A genome-wide mapping of human gene interaction networks revealed also genetic interactions between *XIRP1* and *FGFR1* as wells as *LMBRD1*, which for its part has a close linkage with *LMBR1* over protein domain sharing (IPR006876, PF04791) and *SHH* (Lin *et al.* 2010; Finn *et al.* 2015). The genes *FGFR1, LMBR1* and *SHH* were parts of the candidate gene list associated with polydactyly and syndactyly.

In addition, interactions between *XIRP1, CTNNB1* and *CTNND1* have previously been identified (Wang *et al.* 2010; Wang *et al.* 2013). *CTNNB1* and *CTNND1* affect cadherin function and regulation in cell-to-cell adhesion, motility, and proliferation processes, which is essential for the early embryonic development and the preservation of normal species specific tissue organization (Halbleib and nelson 2006; Wildenberg *et al.* 2006). The congenital malformations in both calves included structures, which are characterized by failed cell-fusion while tissue development, which confirm the expected correlation between the detected variants in *XIRP1* and the present phenotypes due to probably early embryonic interactions with *CTNNB1* and *CTNND1*. Studies of *CTNNB1*/*Wnt* signaling pathways displayed this gene being a regulator in chondrocyte differentiation and synovial joint formation (Guo *et al.* 2004) and required for osteoblast lineage differentiation while skeletal precursors (Day *et al.* 2005; Hill *et al.* 2005). Further, *CTNNB1* operates in *SHH* polarization in hair morphogenesis (Gat *et al.* 1998). Mutations in the *CTNND1* and *cadherin 1* (*CDH1*) are highly associated causing blepharocheilodontic syndrome (BCD) in human (Ghoumid *et al.* 2017). The phenotype of this syndrome included dominant features as eyelid malformations, cleft lip or palate and ectodermal dysplasia. Furthermore, malformation or absence of the thyroid gland, atresia ani, neural tube defects, syndactyly or complex limb reduction defects are reported (Gorlin *et al.* 1976; Weaver *et al.* 2010; Ababneh *et al.* 2014; Ghoumid *et al.* 2017). Both calves showed parts of these phenotypic variations.

Therefore, we propose that we identified a *XIPR1* haplotype containing two de novo variants associated with unilateral polydactyly and polysyndactyly in Holstein cattle.

In conclusion, congenital malformations in cattle have practical relevance and have a high impact on animal welfare. Until now, the genetic background of polydactyly has not been elucidated completely. This makes our study as an important step for understanding the inherited defects and getting the ability for preventing the occurrence of affected animals.

The detection of causal variants for the phenotype polydactyly in cattle is difficult, particularly for sporadic cases. Nevertheless, we suggested an effect of the transformed *XIRP1* protein on the embryonic link between *XIRP1* and the genes *FGFR1, CTNNB1* and *CTNND1*, which are highly associated with diverse phenotypes of isolated unilateral polydactyly and polysyndactyly cases in Holstein cattle, occurring in an incomplete penetrance.

## Acknowledgments

The laboratory work of H. Klippert-Hasberg, N. Wagner and M. Drabert is greatly appreciated. We thank J. Wrede for supporting in computational analysis and Ralph Ingomar for graphical assistance. We gratefully acknowledge support from the North-German Supercomputing Alliance (HLRN) for HPC-resources that contributed to the research results. We are grateful to K. Liebig and H. Lönneker for animal care. For providing sperm samples from the sire bulls, our thank goes to Alta Deutschland GmbH, Uelzen, Germany, and Semex Deutschland GmbH, Verden, Germany.

## Supplemental material

**Figure S1.** Sequence of the bovine ZRS region in *LMBR1* gene matched to Homo sapiens, Mus musculus, Gallus gallus, Canis familiaris and Felis catus. Previously reported variants, which are associated with polydactyly in the different species, are framed. Identical sequences between the species are marked with an asterisk.

**Figure S2.** Sequence of the bovine pZRS region in *LMBR1* gene matched to Homo sapiens, Mus musculus, Canis familiaris and Felis catus. Previously reported variants, which are associated with polydactyly in the different species, are framed. Identical sequences between the species are marked with an asterisk.

**Figure S3.** Gene model of the bovine *LMBR1* gene (ENSBTAG00000030817, Transcript ENSBTAT00000043570.3/ ENSBTAT00000043570.4) located on chromosome 4 according to UMD3.1 and ARS-UCD1.2. The pZRS and ZRS region in intron 5 is marked in bold. For variant analysis by sanger sequencing, three overlapping amplicons were produced. The first location information were according to UMD3.1 and the second in parentheses were accordin to ARS-UCD1.2.

**Figure S4.** Validation results of the familial animals of the two polydactylous calves. (A) edigree of the first female affected calf (case 1) and (B) pedigree of the second male affected calf (case 2). The wild type (wt) and mutant (mut) occurrence of the variants (g.12552417dupC, g.12552582delA) were added to the pedigree for each genotyped animal, were samples were available.

**Table S1.** Candidate genes for the phenotypes polydactyly, syndactyly and polysyndactyly in domestic animals according to NCBI and OMIA. The bovine orthologues genes were presented with their chromosomal position.

**Table S2.** Primer pairs used for validation of the three variants, filtered out from WGS data, *CERK*:g.118035410_128del by PCR amplicon and the two variants *XIRP1*:g.12552417dupC and *XIRP1*:g.12552582delA by Sanger sequencing of the 898 bp amplicon and genotyping by RFLP. Primer pairs sequences, length, annealing temperature (AT), amplicon size (AS) in base pairs (bp) for wild type (wt) and mutant (mut) allele, restriction enzyme, buffer and incubation temperature (IT) are given.

**Table S3.** Primer sequences used for NGS data validation of the ZRS and pZRS region of *LMBR1* gene using Sanger sequencing. Primer pairs, length, annealing temperature (AT) and amplicon size (AS) in base pairs (bp) and are given.

**Table S4.** Results of the filtered whole genome sequencing data. All variants, which are homozygous or heterozygous mutant exclusively in the affected polydactylous calves, combined with heterozygous mutant or homozygous wild type alleles in the dam of Case 2, were chosen. The remaining critical variants for the phenotype polydactyly and polysyndactyly, which were validated, are printed in bold.

**Table S5.** Genotyping results of the filtered variants (g.12552417dupC, g.12552582delA) in *XIRP1* in 359 cattle of 8 different breeds using restriction fragment length polymorphisms (RFLP). The distribution of the wild type allele (wt) and mutant allele (mut) is shown. Seven animals of the Holstein breed were heterozygous and one homozygous carrier of the combination of the variants g.12552417dupC and g.12552582delA. None of the variants were detected in the other breeds.

## REFERENCES

Ababneh, F. K., A. Al Swaid, A. Elhag, T. Youssef and S. Alsaif, 2014 Blepharo-cheilo-dontic (BCD) syndrome: Expanding the phenotype, case report and review of literature. American Journal of Medical Genetics Part A 164: 1525–1529.

Acuna-Hidalgo, R., T. Bo, M. P. Kwint, M. Van De Vorst, M. Pinelli et al., 2015 Post-zygotic point mutations are an underrecognized source of de novo genomic variation. The American Journal of Human Genetics 97: 67–74.

Al Qattan, M. M., I. Al Abdulkareem, Y. Al Haidan and M. Al Balwi, 2012 A novel mutation in the SHH long-range regulator (ZRS) is associated with preaxial polydactyly, triphalangeal thumb, and severe radial ray deficiency. American Journal of Medical Genetics Part A 158: 2610–2615.

Alam, M., J. Lee, H. Lee, J. Ko, K. Lee et al., 2007 Supernumerary ectopic limbs in Korean indigenous cattle: four case reports. Veterinarni Medicina-Praha- 52: 202.

Albuisson, J., B. Isidor, M. Giraud, O. Pichon, T. Marsaud et al., 2011 Identification of two novel mutations in Shh long-range regulator associated with familial pre-axial polydactyly. Clinical genetics 79: 371–377.

Altenbrunner-Martinek, B., E. Gruber and W. Baumgartner, 2007 Fallbeschreibung: Polysyndaktylie bei einem osterreichischen Fleckvieh-Stierkalb. Wiener Tieraerztliche Monatsschrift 94: 287.

Andrews, S., 2015 A quality control tool for high throughput sequence data. 2010. Google Scholar.

Bähr, C., K. Wittenberg and O. Distl, 2003 Case report: polydactyly in a German holstein calf. DTW. Deutsche tierarztliche Wochenschrift 110: 333–335.

Campbell, I. M., B. Yuan, C. Robberecht, R. Pfundt, P. Szafranski et al., 2014 Parental somatic mosaicism is underrecognized and influences recurrence risk of genomic disorders. The American Journal of Human Genetics 95: 173–182.

Carstanjen, B., J. Pennecke, S. Boehart and K. E. Müller, 2010 Unilateral polydactylism in a German Holstein-Friesian calf-A case report. The Thai Journal of Veterinary Medicine 40: 69–74.

Cingolani, P., A. Platts, L. L. Wang, M. Coon, T. Nguyen et al., 2012 A program for annotating and predicting the effects of single nucleotide polymorphisms, SnpEff: SNPs in the genome of Drosophila melanogaster strain w1118; iso-2; iso-3. Fly 6: 80– 92.

Credille, K., K. Barnhart, J. Minor and R. Dunstan, 2005 Mild recessive epidermolytic hyperkeratosis associated with a novel keratin 10 donor splice-site mutation in a family of Norfolk terrier dogs. British Journal of Dermatology 153: 51–58.

Day, T. F., X. Guo, L. Garrett-Beal and Y. Yang, 2005 Wnt/β-catenin signaling in mesenchymal progenitors controls osteoblast and chondrocyte differentiation during vertebrate skeletogenesis. Developmental cell 8: 739–750.

Dorshorst, B., R. Okimoto and C. Ashwell, 2010 Genomic regions associated with dermal hyperpigmentation, polydactyly and other morphological traits in the Silkie chicken. Journal of Heredity 101: 339–350.

Dunn, I. C., I. R. Paton, A. K. Clelland, S. Sebastian, E. J. Johnson et al., 2011 The chicken polydactyly (Po) locus causes allelic imbalance and ectopic expression of Shh during limb development. Developmental Dynamics 240: 1163–1172.

Farooq, M., J. T. Troelsen, M. Boyd, H. Eiberg, L. Hansen et al., 2010 Preaxial polydactyly/triphalangeal thumb is associated with changed transcription factor-binding affinity in a family with a novel point mutation in the long-range cisregulatory element ZRS. European Journal of Human Genetics 18: 733.

Feng, H.-Z., Q. Wang, R. S. Reiter, J. L.-C. Lin, J. J.-C. Lin et al., 2013 Localization and function of Xinα in mouse skeletal muscle. American Journal of Physiology-Cell Physiology 304: C1002–C1012.

Finn, R. D., P. Coggill, R. Y. Eberhardt, S. R. Eddy, J. Mistry et al., 2015 The Pfam protein families database: towards a more sustainable future. Nucleic acids research 44: D279–D285.

Freick, M., H. Behn, M. Hardt, S. Knobloch, B. Eftekharzadeh et al., 2014 Monozygotic incomplete caudal duplication in a German Holstein calf. Veterinary Record Case Reports 2: e000048.

Furniss, D., L. A. Lettice, I. B. Taylor, P. S. Critchley, H. Giele et al., 2008 A variant in the sonic hedgehog regulatory sequence (ZRS) is associated with triphalangeal thumb and deregulates expression in the developing limb. Human molecular genetics 17: 2417– 2423.

Gat, U., R. DasGupta, L. Degenstein and E. Fuchs, 1998 De novo hair follicle morphogenesis and hair tumors in mice expressing a truncated β-catenin in skin. Cell 95: 605–614.

Ghoumid, J., M. Stichelbout, A.-S. Jourdain, F. Frenois, S. Lejeune-Dumoulin et al., 2017 Blepharocheilodontic syndrome is a CDH1 pathway–related disorder due to mutations in CDH1 and CTNND1. Genetics in Medicine 19: 1013.

Girisha, K. M., A. M. Bidchol, P. S. Kamath, K. H. Shah, G. R. Mortier et al., 2014 A novel mutation (g. 106737G> T) in zone of polarizing activity regulatory sequence (ZRS) causes variable limb phenotypes in Werner mesomelia. American Journal of Medical Genetics Part A 164: 898–906.

Gorlin, R., J. Pindborg and M. Cohen, 1976 Hemifacial atrophy. Syndromes of the head and neck. New York: McGraw-Hill: 546–552.

Grosskurth, S. E., D. Bhattacharya, Q. Wang and J. J.-C. Lin, 2008 Emergence of Xin demarcates a key innovation in heart evolution. PLoS One 3: e2857.

Guerrero Netro, H. M., 2014 Differential regulation of early response genes by fibroblast growth factor (FGF) 8 and FGF18 in bovine granulosa cells in vitro.

Gugjoo, M. B., I. P. Sarode and S. Kumar, 2013 Bilateral Polydactyly in a Nondescript Calf. Journal of Advanced Veterinary Research 3: 86–88.

Guo, X., T. F. Day, X. Jiang, L. Garrett-Beal, L. Topol et al., 2004 Wnt/β-catenin signaling is sufficient and necessary for synovial joint formation. Genes & development 18: 2404– 2417.

Gurnett, C. A., A. M. Bowcock, F. R. Dietz, J. A. Morcuende, J. C. Murray et al., 2007 Two novel point mutations in the long-range SHH enhancer in three families with triphalangeal thumb and preaxial polydactyly. American Journal of Medical Genetics Part A 143: 27–32.

Hajihosseini, M. K., M. D. Lalioti, S. Arthaud, H. R. Burgar, J. M. Brown et al., 2004 Skeletal development is regulated by fibroblast growth factor receptor 1 signalling dynamics. Development 131: 325–335.

Halbleib, J. M., and W. J. Nelson, 2006 Cadherins in development: cell adhesion, sorting, and tissue morphogenesis. Genes & development 20: 3199–3214.

Hill, T. P., D. Später, M. M. Taketo, W. Birchmeier and C. Hartmann, 2005 Canonical Wnt/βcatenin signaling prevents osteoblasts from differentiating into chondrocytes. Developmental cell 8: 727–738.

Hovius, S. E., 2018 A point mutation in the pre-ZRS disrupts sonic hedgehog expression in the limb bud and results in triphalangeal thumb–polysyndactyly syndrome. Genet Med.

Huang, Y. Q., X. M. Deng, Z. Q. Du, X. Qiu, X. Du et al., 2006 Single nucleotide polymorphisms in the chicken Lmbr1 gene are associated with chicken polydactyly. Gene 374: 10–18.

Ingham, P. W., and A. P. McMahon, 2001 Hedgehog signaling in animal development: paradigms and principles. Genes & development 15: 3059–3087.

Johnson, E. J., D. M. Neely, I. C. Dunn and M. G. Davey, 2014 Direct functional consequences of ZRS enhancer mutation combine with secondary long range SHH signalling effects to cause preaxial polydactyly. Developmental biology 392: 209–220.

Johnson, J., H. Leipold, M. Guffy, S. Dennis and R. Schalles, 1982 Characterization of bovine polydactyly [Anatomy, genetics]. Bovine practice.

Johnson, J., H. Leipold, R. Schalles, M. Guffy, J. Peeples et al., 1981 Hereditary polydactyly in Simmental cattle. Journal of heredity 72: 205–208.

Kaus, W., 1968 Syn-polydactyly in an ox. Deutsche tierärztliche Wochenschrift 75: 120.

Kim, C.-s., S.-c. Yeon, G.-h. Cho, J.-h. Lee, M.-c. Choi et al., 2001 Polymelia with two extra forelimbs at the right scapular region in a male Korean native calf. Journal of Veterinary Medical Science 63: 1161–1164.

Krugljak, T., 2014 Genealogical kinship Holstein bulls in the bloodlines. Науковий вісник НУБіП України. Серія: Технологія виробництва і переробки продукції тваринництва..

Larkin, M. A., G. Blackshields, N. Brown, R. Chenna, P. A. McGettigan et al., 2007 Clustal W and Clustal X version 2.0. bioinformatics 23: 2947–2948.

Laurell, T., J. E. VanderMeer, A. M. Wenger, G. Grigelioniene, A. Nordenskjöld et al., 2012 A novel 13 base pair insertion in the sonic hedgehog ZRS limb enhancer (ZRS/LMBR1) causes preaxial polydactyly with triphalangeal thumb. Human mutation 33: 1063–1066.

Lettice, L. A., S. J. Heaney, L. A. Purdie, L. Li, P. de Beer et al., 2003 A long-range Shh enhancer regulates expression in the developing limb and fin and is associated with preaxial polydactyly. Human molecular genetics 12: 1725–1735.

Lettice, L. A., A. E. Hill, P. S. Devenney and R. E. Hill, 2008 Point mutations in a distant sonic hedgehog cis-regulator generate a variable regulatory output responsible for preaxial polydactyly. Human molecular genetics 17: 978–985.

Li, H., and R. Durbin, 2009 Fast and accurate short read alignment with Burrows–Wheeler transform. Bioinformatics 25: 1754–1760.

Li, H., B. Handsaker, A. Wysoker, T. Fennell, J. Ruan et al., 2009 The sequence alignment/map format and SAMtools. Bioinformatics 25: 2078–2079.

Lin, A., R. T. Wang, S. Ahn, C. C. Park and D. J. Smith, 2010 A genome-wide map of human genetic interactions inferred from radiation hybrid genotypes. Genome research.

Maas, S. A., T. Suzuki and J. F. Fallon, 2011 Identification of spontaneous mutations within the long-range limb-specific Sonic hedgehog enhancer (ZRS) that alter Sonic hedgehog expression in the chicken limb mutants oligozeugodactyly and silkie breed. Developmental Dynamics 240: 1212–1222.

Manokaran, S., M. Palanisamy, S. Prakash, M. Selvaraju and R. E. Napoleon, 2018 Unilateral Polydactylism with Brachygnathism in a Calf.

Marolt, J., and B. Ilijas, 1967 Contribution to polydactyly in cattle. Deutsche tierärztliche Wochenschrift 74: 197.

Masuya, H., H. Sezutsu, Y. Sakuraba, T. Sagai, M. Hosoya et al., 2007 A series of ENU-induced single-base substitutions in a long-range cis-element altering Sonic hedgehog expression in the developing mouse limb bud. Genomics 89: 207–214.

Mather, D., 1987 Polydactyly in calves. Veterinary Record 120: 487–487.

McKenna, A., M. Hanna, E. Banks, A. Sivachenko, K. Cibulskis et al., 2010 The Genome Analysis Toolkit: a MapReduce framework for analyzing next-generation DNA sequencing data. Genome research 20: 1297–1303.

Mineyko, A., A. Doja, J. Hurteau, W. B. Dobyns, S. Das et al., 2010 A novel missense mutation in LIS1 in a child with subcortical band heterotopia and pachygyria inherited from his mildly affected mother with somatic mosaicism. Journal of child neurology 25: 738–741.

Morrill, E., 1945 A new sex-linked defect in cattle. Journal of Heredity 36: 81-82.

Murondoti, A., and R. Busayi, 2001 Perineomelia, polydactyly and other malformations in a Mashona calf. Veterinary Record 148: 512–513.

Norbnop, P., C. Srichomthong, K. Suphapeetiporn and V. Shotelersuk, 2014 ZRS 406A> G mutation in patients with tibial hypoplasia, polydactyly and triphalangeal first fingers. Journal of human genetics 59: 467.

Ojo, S., H. Leipold, G. Saperstein and M. Guffy, 1975 Polydactyly in a Holstein-Friesian calf. Giessener Beitraege zur Erbpathologie und Zuchthygiene (Germany, FR).

Park, K., J. Kang, K. P. Subedi, J.-H. Ha and C. Park, 2008 Canine polydactyl mutations with heterogeneous origin in the conserved intronic sequence of LMBR1. Genetics 179: 2163–2172.

Rafee, M., and D. Madhu, 2016 Surgical Management of Bilateral Polydactyly in Twin Calves. SKUAST Journal of Research 18: 62–64.

Roberts, E., 1921 Polydactylism in cattle. Journal of Heredity 12: 84–86.

Sangwan, V., S. Mahajan, A. Kumar, A. Anand and N. Saini, 2015 Unilateral Polydactyly in a Holstein Friesian Heifer. Indian Vet. J 92: 52–53.

Schmidt, T. A., J. Wrede and D. L. Simon, 2006 OPTI-MATE Ver. 4.0, Management-Computer-Programm zur Minimierung der Inzucht in gefährdeten Populationen.

Schmieder, R., and R. Edwards, 2011 Quality control and preprocessing of metagenomic datasets. Bioinformatics 27: 863–864.

Schönfelder, A., T. Wittek and A. Sobiraj, 2003 Die Polymelie beim Kalb-Übersicht mit Fallbeschreibungen zur operativen Behebung. Tierärztliche Praxis Großtiere 31: 314– 318.

Schummer, A., 1935 Entwicklungsbedingte Polydaktylic beim Rinde. Zeitschrift für Anatomie und Entwicklungsgeschichte 104: 491–501.

Semerci, C., F. Demirkan, M. Özdemir, E. Biskin, B. Akin et al., 2009 Homozygous feature of isolated triphalangeal thumb–preaxial polydactyly linked to 7q36: no phenotypic difference between homozygotes and heterozygotes. Clinical genetics 76: 85–90.

Sicca, F., A. Kelemen, P. Genton, S. Das, D. Mei et al., 2003 Mosaic mutations of the LIS1 gene cause subcortical band heterotopia. Neurology 61: 1042–1046.

Sindi, S. S., S. Önal, L. C. Peng, H.-T. Wu and B. J. Raphael, 2012 An integrative probabilistic model for identification of structural variation in sequencing data. Genome biology 13: R22.

Spadari, A., G. Spinella, A. Venturini and A. Gentile, 2003 Fore limb bilateral polydactyly and ocular dermoid in a Holstein Friesian calf. Atti della Societa italiana di buiatria 35: 149–155.

Thisse, B., and C. Thisse, 2005 Functions and regulations of fibroblast growth factor signaling during embryonic development. Developmental biology 287: 390–402.

Tyson, R., J. P. Graham, P. T. Colahan and C. R. Berry, 2004 Skeletal atavism in a miniature horse. Veterinary Radiology & Ultrasound 45: 315–317.

VanderMeer, J. E., M. Afzal, S. Alyas, S. Haque, N. Ahituv et al., 2012 A novel ZRS mutation in a Balochi tribal family with triphalangeal thumb, pre-axial polydactyly, post-axial polydactyly, and syndactyly. American Journal of Medical Genetics Part A 158: 2031–2035.

VanderMeer, J. E., and N. Ahituv, 2011 cis-regulatory mutations are a genetic cause of human limb malformations. Developmental Dynamics 240: 920–930.

VanderMeer, J. E., R. Lozano, M. Sun, Y. Xue, D. Daentl et al., 2014 A novel ZRS mutation leads to preaxial polydactyly type 2 in a heterozygous form and Werner mesomelic syndrome in a homozygous form. Human mutation 35: 945–948.

Vermunt, J., H. Burbidge and K. Thompson, 2000 Unusual congenital deformities of the lower limb in two calves. New Zealand veterinary journal 48: 192–194.

Wang, Q., J. L.-C. Lin, B. E. Reinking, H.-Z. Feng, F.-C. Chan et al., 2010 Essential roles of an intercalated disc protein, mXinβ, in postnatal heart growth and survival. Circulation research 106: 1468.

Wang, Q., T.-L. Lu, E. Adams, J. L.-C. Lin and J. J.-C. Lin, 2013 Intercalated disc protein, mXinα, suppresses p120-catenin-induced branching phenotype via its interactions with p120-catenin and cortactin. Archives of biochemistry and biophysics 535: 91– 100.

Warde-Farley, D., S. L. Donaldson, O. Comes, K. Zuberi, R. Badrawi et al., 2010 The GeneMANIA prediction server: biological network integration for gene prioritization and predicting gene function. Nucleic acids research 38: W214–W220.

Weaver, K. N., K. D. Rutledge, J. H. Grant and N. H. Robin, 2010 Imperforate anus is a rare associated finding in blepharocheilodontic syndrome. American Journal of Medical Genetics Part A 152: 438–440.

Wieczorek, D., B. Pawlik, Y. Li, N. A. Akarsu, A. Caliebe et al., 2009 A specific mutation in the distant sonic hedgehog (SHH) cis-regulator (ZRS) causes Werner mesomelic syndrome (WMS) while complete ZRS duplications underlie Haas type polysyndactyly and preaxial polydactyly (PPD) with or without triphalangeal thumb. Human mutation 31: 81–89.

Wildenberg, G. A., M. R. Dohn, R. H. Carnahan, M. A. Davis, N. A. Lobdell et al., 2006 p120-catenin and p190RhoGAP regulate cell-cell adhesion by coordinating antagonism between Rac and Rho. Cell 127: 1027–1039.

Wockl, F., B. Mayr and W. Schleger, 1980 Moglichkeiten der Diagnose von Erbfehlern mit Hilfe der Chromosomenanalyse bei Rind, Pferd und Schwein--eine Ubersicht. Berliner und Munchener tierarztliche Wochenschrift.

Wu, P.-F., S. Guo, X.-F. Fan, L.-L. Fan, J.-Y. Jin et al., 2016 A novel ZRS mutation in a Chinese patient with Preaxial Polydactyly and Triphalangeal thumb. Cytogenetic and genome research 149: 171–175.

Xiang, Y., L. Jiang, B. Wang, Y. Xu, H. Cai et al., 2017 Mutational screening of GLI3, SHH, preZRS, and ZRS in 102 Chinese children with nonsyndromic polydactyly. Developmental Dynamics.

