## Supplementary Tables 1-5 for "*De-novo* variants in *XIRP1* associated with polydactyly and polysyndactyly in Holstein cattle"

**Table S1.** Candidate genes for the phenotypes polydactyly, syndactyly and polysyndactyly in domestic animals according to NCBI and OMIA. The bovine orthologues genes were presented with their chromosomal position.

| **Phenotype** | **Gene** | **Gene full name** | **Species** | **Gene ID** | **Bovine gene** | **BTA** | **Position** |
|---|---|---|---|---|---|---|---|
| Syndactyly | *ADAMTS5* | a disintegrin-like and metallopeptidase (reprolysin type) with thrombospondin type 1 motif, 5 (aggrecanase-2) | Mus musculus | ENSBTAG00000000648 | *ADAMTS5* | 1 | 8811792-8861602 |
| Polydactyly | *ARL13B* | ADP ribosylation factor like GTPase 13B | Homo sapiens | ENSBTAG00000005155 | *ARL13B* | 1 | 37875395-37952674 |
| Polydactyly | *ARL6* | ADP ribosylation factor like GTPase 6 | Homo sapiens | ENSBTAG00000010091 | *ARL6* | 1 | 41640353-41679711 |
| Polydactyly | *Dzip1l* | DAZ interacting protein 1-like | Mus musculus | ENSBTAG00000015198 | *DZIP1L* | 1 | 132103024-132129289 |
| Polydactyly | *Etv5* | ets variant 5 | Mus musculus | ENSBTAG00000014915 | *ETV5* | 1 | 81725703-81784999 |
| Polydactyly | *IFT80* | intraflagellar transport 80 | Homo sapiens | ENSBTAG00000031572 | *IFT80* | 1 | 107962966-108058677 |
| Polydactyly, syndactyly | *PIK3CA* | phosphatidylinositol-4,5-bisphosphate 3-kinase catalytic subunit alpha | Homo sapiens | ENSBTAG00000009232 | *PIK3CA* | 1 | 88504290-88533061 |
| Polydactyly | *TCTEX1D2* | Tctex1 domain containing 2 | Homo sapiens | ENSBTAG00000032684 | *TCTEX1D2* | 1 | 71489285-71516789 |
| Polydactyly | *BBS5* | Bardet-Biedl syndrome 5 | Homo sapiens | ENSBTAG00000010736 | *BBS5* | 2 | 26800661-26821659 |
| Polydactyly | *CCDC28B* | coiled-coil domain containing 28B | Homo sapiens | ENSBTAG00000015104 | *CCDC28B* | 2 | 122150084-122154154 |
| Polydactyly, syndactyly | *GLI2* | GLI family zinc finger 2 | Homo sapiens | ENSBTAG00000011682 | *GLI2* | 2 | 72977209-73168370 |
| Polydactyly | *Hoxd9* | homeobox D9 | Mus musculus | ENSBTAG00000016033 | *HOXD9* | 2 | 20824235-20825910 |
| Polydactyly | *Hoxd10* | homeobox D10 | Mus musculus | ENSBTAG00000016030 | *HOXD10* | 2 | 20828934-20832040 |
| Polydactyly, syndactyly | *Hoxd13* | homeobox D13 | Mus musculus | ENSBTAG00000004313 | *HOXD13* | 2 | 20854099-20856159 |
| Polydactyly, syndactyly | *Ihh* | Indian hedgehog | Mus musculus | ENSBTAG00000008452 | *IHH* | 2 | 107722668-107728939 |
| Polydactyly | *PDE6D* | phosphodiesterase 6D | Homo sapiens | ENSBTAG00000019480 | *PDE6D* | 2 | 120250522-120305878 |
| Polydactyly | *TMEM237* | transmembrane protein 237 | Homo sapiens | ENSBTAG00000017437 | *TMEM237* | 2 | 90588104-90610465 |

| **Phenotype** | **Gene** | **Gene full name** | **Species** | **Gene ID** | **Bovine gene** | **BTA** | **Position** |
|---|---|---|---|---|---|---|---|
| Polydactyly | *TTC21B* | tetratricopeptide repeat domain 21B | Homo sapiens | ENSBTAG00000016512 | *TTC21B* | 2 | 30433135-30531188 |
| Syndactyly | *NECTIN4* | nectin cell adhesion molecule 4 | Homo sapiens | ENSBTAG00000017877 | *NECTIN4* | 3 | 8421828-8438222 |
| Polydactyly | *Traf3ip1* | TRAF3 interacting protein 1 | Mus musculus | ENSBTAG00000015973 | *TRAF3IP1* | 3 | 118273223-118334789 |
| Polydactyly | *BBS9* | Bardet-Biedl syndrome 9 | Homo sapiens | ENSBTAG00000006528 | *BBS9* | 4 | 63649808-64105542 |
| Polydactyly | *CEP41* | centrosomal protein 41 | Homo sapiens | ENSBTAG00000007413 | *CEP41* | 4 | 94959111-95006794 |
| Polydactyly, polysyndactyly | *Gli3* | GLI-Kruppel family member GLI3 | Homo sapiens Mus musculus | ENSBTAG00000010671 | *GLI3* | 4 | 79444243-79758476 |
| Polydactyly | *Hdac9* | histone deacetylase 9 | Mus musculus | ENSBTAG00000003808 | *HDAC9* | 4 | 27174250-27457097 |
| Polydactyly | *Hoxa11* | homeobox A11 | Mus musculus | ENSBTAG00000014738 | *HOXA11* | 4 | 69311987-69314613 |
| Polydactyly | *HOXA13* | homeobox A13 | Homo sapiens | ENSBTAG00000014735 | *HOXA13* | 4 | 69297275-69298937 |
| Polydactyly, syndactyly, olysyndactyly | *LMBR1* | limb development membrane protein 1 | Homo sapiens Mus musculus Gallus gallus | ENSBTAG00000030817 | *LMBR1* | 4 | 118948100-119080458 |
| Polydactyly, syndactyly, polysyndactyly | *SHH* | sonic hedgehog | Homo sapiens Mus musculus Felis catus Gallus gallus | ENSBTAG00000024552 | *SHH* | 4 | 118265405-118274566 |
| Polysyndactyly | *SMO* | Smoothened, frizzled class receptor | Homo sapdiens | ENSBTAG00000013287 | *SMO* | 4 | 93922581-93946896 |
| Polydactyly, syndactyly | *Twist1* | twist basic helix-loop-helix transcription factor 1 | Mus musculus | ENSBTAG00000046922 | *TWIST1* | 4 | 27854574-27855179 |
| Polydactyly | *WDR60* | WD repeat domain 60 | Homo sapiens | ENSBTAG00000024157 | *WDR60* | 4 | 120326813-120373331 |
| Syndactyly | *ADAMTS20* | a disintegrin-like and metallopeptidase (reprolysin type) with thrombospondin type 1 motif, 20 | Mus musculus | ENSBTAG00000000395 | *ADAMTS20* | 5 | 37102590-37332440 |
| Polydactyly | *BBS10* | Bardet-Biedl syndrome 10 | Homo sapiens | ENSBTAG00000004425 | *BBS10* | 5 | 5813257-5816003 |
| Syndactyly | *CACNA1C* | calcium voltage-gated channel subunit alpha1 C | Homo sapiens | ENSBTAG00000010660 | *CACNA1C* | 5 | 109152548-109417890 |
| Polydactyly | *CCND2* | cyclin D2 | Homo sapiens | ENSBTAG00000016649 | *CCND2* | 5 | 106253907-106276819 |

| Phenotype | Gene | Gene full name | Species | Gene ID | Bovine gene | BTA | Position |
|---|---|---|---|---|---|---|---|
| Polydactyly | *CEP290* | centrosomal protein 290 | Homo sapiens | ENSBTAG00000018745 | *CEP290* | 5 | 17902277-17998418 |
| Polydactyly | *BBS7* | Bardet-Biedl syndrome 7 | Homo sapiens | ENSBTAG00000004945 | *BBS7* | 6 | 3375322-3410957 |
| Polydactyly | *CC2D2A* | coiled-coil and C2 domain containing 2A | Homo sapiens | ENSBTAG00000018638 | *CC2D2A* | 6 | 115477255-115563715 |
| Polydactyly | *EVC* | EvC ciliary complex subunit 1 | Homo sapiens | ENSBTAG00000004287 | *EVC* | 6 | 105171716-105275204 |
| Polydactyly | *EVC2* | EvC ciliary complex subunit 2 | Homo sapiens | ENSBTAG00000004277 | *EVC2* | 6 | 105291556-105452049 |
| Syndactyly | *FRAS1* | Fraser extracellular matrix complex subunit 1 | Mus musculus | ENSBTAG00000010716 | *FRAS1* | 6 | 94711652-95055569 |
| Polydactyly | *MSX1* | msh homeobox 1 | Homo sapiens | ENSBTAG00000010875 | *MSX1* | 6 | 106061463-106065759 |
| Polydactyly | *WDR19* | WD repeat domain 19 | Homo sapiens | ENSBTAG00000014512 | *WDR19* | 6 | 59993685-60065797 |
| Polydactyly | *CEP120* | centrosomal protein 120 | Homo sapiens | ENSBTAG00000013184 | *CEP120* | 7 | 31688655-31774451 |
| Polydactyly | *PIK3R2* | phosphoinositide-3-kinase regulatory subunit 2 | Homo sapiens | ENSBTAG00000002350 | *PIK3R2* | 7 | 4987071-4999686 |
| Polydactyly | *PITX1* | paired like homeodomain 1 | Homo sapiens | ENSBTAG00000004602 | *PITX1* | 7 | 48063871-48069063 |
| Syndactyly | *SMAD5* | SMAD family member 5 | Mus musculus | ENSBTAG00000021772 | *SMAD5* | 7 | 49155483-49217780 |
| Syndactyly | *ESCO2* | establishment of sister chromatid cohesion N-acetyltransferase 2 | Homo sapiens | ENSBTAG00000006551 | *ESCO2* | 8 | 10859329-10885722 |
| Polydactyly | *Hand2* | heart and neural crest derivatives expressed 2 | Mus musculus | ENSBTAG00000044123 | *HAND2* | 8 | 5747710-5749915 |
| Polydactyly | *NEK1* | NIMA related kinase 1 | Homo sapiens | ENSBTAG00000026915 | *NEK1* | 8 | 1266978-1394750 |
| Syndactyly | *ROR2* | receptor tyrosine kinase like orphan receptor 2 | Homo sapiens | ENSBTAG00000005092 | *ROR2* | 8 | 87340388-87578642 |
| Polydactyly | *TRIM32* | tripartite motif containing 32 | Homo sapiens | ENSBTAG00000017155 | *TRIM32* | 8 | 107645159-107656813 |
| Polydactyly | *AHI1* | Abelson helper integration site 1 | Homo sapiens | ENSBTAG00000017958 | *AHI1* | 9 | 74330947-74544060 |
| Syndactyly | *GJA1* | gap junction protein alpha 1 | Homo sapiens | ENSBTAG00000001835 | *GJA1* | 9 | 30127786-30140793 |
| Polydactyly | *BBS4* | Bardet-Biedl syndrome 4 | Homo sapiens | ENSBTAG00000007614 | *BBS4* | 10 | 19317484-19363592 |
| Polydactyly | *IFT43* | intraflagellar transport 43 | Homo sapiens | ENSBTAG00000012005 | *IFT43* | 10 | 88272302-88379899 |
| Polydactyly | *KIAA0586* | KIAA0586 | Homo sapiens | ENSBTAG00000004631 | *KIAA0586* | 10 | 70945020-71078172 |
| Polydactyly | *TTC8* | tetratricopeptide repeat domain 8 | Homo sapiens | ENSBTAG00000030432 | *TTC8* | 10 | 101571088-101629775 |

| Phenotype | Gene | Gene full name | Species | Gene ID | Bovine gene | BTA | Position |
|---|---|---|---|---|---|---|---|
| Syndactyly | *CKAP2L* | cytoskeleton associated protein 2 like | Homo sapiens | ENSBTAG00000002298 | *CKAP2L* | 11 | 46318613-46340841 |
| Polydactyly | *DYNC2LI1* | dynein cytoplasmic 2 light intermediate chain 1 | Homo sapiens | ENSBTAG00000013676 | *DYNC2LI1* | 11 | 26077551-26115657 |
| Polydactyly | *IFT172* | intraflagellar transport 172 | Homo sapiens | ENSBTAG00000018157 | *IFT172* | 11 | 72185482-72222959 |
| Polydactyly | *INPP5E* | inositol polyphosphate-5-phosphatase E | Homo sapiens | ENSBTAG00000001354 | *INPP5E* | 11 | 103928312-103936269 |
| Syndactyly | *MYCN* | MYCN proto-oncogene, bHLH transcription factor | Homo sapiens<br>Mus musculus | ENSBTAG00000023372 | N/A | 11 | 82694507-82697306 |
| Polydactyly | *NPHP1* | nephrocystin 1 | Homo sapiens | ENSBTAG00000007675 | *NPHP1* | 11 | 1670784-1735642 |
| Polydactyly | *WDPCP* | WD repeat containing planar cell polarity effector | Homo sapiens | ENSBTAG00000005151 | *WDPCP* | 11 | 61759950-61969748 |
| Polydactyly | *WDR34* | WD repeat domain 34 | Homo sapiens | ENSBTAG00000015343 | *WDR34* | 11 | 99179458-99198829 |
| Polydactyly | *IFT52* | intraflagellar transport 52 | Homo sapiens | ENSBTAG00000019606 | *IFT52* | 13 | 72886683-72924464 |
| Polydactyly | *MKKS* | McKusick-Kaufman syndrome | Homo sapiens | ENSBTAG00000034987 | *MKKS* | 13 | 3574758-3582527 |
| Polydactyly | *SALL4* | spalt like transcription factor 4 | Homo sapiens | ENSBTAG00000003101 | *SALL4* | 13 | 80297262-80313158 |
| Polydactyly | *C8orf37* | chromosome 8 open reading frame 37 | Homo sapiens | ENSBTAG00000043970 | *C8ORF37* | 14 | 71396805-71419794 |
| Polydactyly | *CSPP1* | centrosome and spindle pole associated protein 1 | Homo sapiens | ENSBTAG00000014689 | *CSPP1* | 14 | 33243588-33342863 |
| Polydactyly | *IFT27* | intraflagellar transport 27 | Homo sapiens | ENSBTAG00000006213 | N/A | 14 | 58850048-58851086 |
| Polydactyly | *TMEM67* | transmembrane protein 67 | Homo sapiens | ENSBTAG00000044190 | N/A | 14 | 72788487-72816854 |
| Polydactyly, syndactyly | *Alx4* | aristaless-like homeobox 4 | Mus musculus | ENSBTAG00000027563 | *ALX4* | 15 | 75154393-75187019 |
| Syndactyly | *LRP4* | LDL receptor related protein 4 | Homo sapiens<br>Bos taurus | ENSBTAG00000008429 | *LRP4* | 15 | 77663792-77701236 |
| Polydactyly | *Zbtb16* | zinc finger and BTB domain containing 16 | Homo sapiens | ENSBTAG00000011266 | *ZBTB16* | 15 | 24917133-25116350 |
| Polydactyly | *AKT3* | AKT serine/threonine kinase 3 | Homo sapiens | ENSBTAG00000017788 | *AKT3* | 16 | 34132648-34404652 |
| Polydactyly | *CEP104* | centrosomal protein 104 | Homo sapiens | ENSBTAG00000020014 | *CEP104* | 16 | 50498425-50533080 |

| Phenotype | Gene | Gene full name | Species | Gene ID | Bovine gene | BTA | Position |
|---|---|---|---|---|---|---|---|
| Syndactyly | *IRF6* | interferon regulatory factor 6 | Homo sapiens | ENSBTAG00000002849 | *IRF6* | 16 | 75401221-75417466 |
| Polydactyly | *SDCCAG8* | serologically defined colon cancer antigen 8 | Homo sapiens | ENSBTAG00000017775 | *SDCCAG8* | 16 | 34415701-34655651 |
| Polydactyly | *BBS12* | Bardet-Biedl syndrome 12 | Homo sapiens | ENSBTAG00000007564 | *BBS12* | 17 | 35334854-35336980 |
| Polydactyly | *DGCR2* | DiGeorge syndrome critical region gene 2 | Homo sapiens | ENSBTAG00000000429 | *DGCR2* | 17 | 74581054-74598153 |
| Polydactyly | *DGCR6* | DiGeorge syndrome critical region gene 6 | Homo sapiens | ENSBTAG00000047299 | *DGCR6L* | 17 | 74049772-74053881 |
| Polydactyly | *DGCR8* | DGCR8, microprocessor complex subunit | Homo sapiens | ENSBTAG00000019869 | *DGCR8* | 17 | 74987854-75002264 |
| Polydactyly | *ESS2* | ess-2 splicing factor homolog | Homo sapiens | ENSBTAG00000018534 | *ESS2* | 17 | 74613480-74620081 |
| Polydactyly | *IFT81* | intraflagellar transport 81 | Homo sapiens | ENSBTAG00000020584 | *IFT81* | 17 | 56311086-56408960 |
| Polydactyly | *INTU* | inturned planar cell polarity protein | Homo sapiens<br>Mus musculus | ENSBTAG00000012824 | *INTU* | 17 | 30324842-30404702 |
| Syndactyly | *SFRP2* | secreted frizzled-related protein 2 | Mus musculus | ENSBTAG00000018563 | *SFRP2* | 17 | 3829564-3838136 |
| Syndactyly | *SMAD1* | SMAD family member 1 | Mus musculus | ENSBTAG00000002835 | *SMAD1* | 17 | 12906376-12953417 |
| Polydactyly | *SPECC1L* | sperm antigen with calponin homology and coiled-coil domains 1 like | Homo sapiens | ENSBTAG00000021656 | *SPECC1L* | 17 | 73595800-73658019 |
| Polydactyly | *TCTN1* | tectonic family member 1 | Homo sapiens | ENSBTAG00000009208 | *TCTN1* | 17 | 56724729-56752196 |
| Polydactyly | *TCTN2* | tectonic family member 2 | Homo sapiens | ENSBTAG00000007271 | *TCTN2* | 17 | 54245927-54274411 |
| Polydactyly | *BBS2* | Bardet-Biedl syndrome 2 | Homo sapiens | ENSBTAG00000014923 | *BBS2* | 18 | 24205242-24231176 |
| Polydactyly | *RPGRIP1L* | RPGRIP1 like | Homo sapiens | ENSBTAG00000012499 | *RPGRIP1L* | 18 | 22021983-22114274 |
| Polydactyly | *SALL1* | spalt like transcription factor 1 | Homo sapiens | ENSBTAG00000008544 | *SALL1* | 18 | 19640447-19654562 |
| Polydactyly | *ZNF141* | zinc finger protein 141 | Homo sapiens | ENSBTAG00000045581<br>ENSBTAG00000030444<br>ENSBTAG00000037699<br>ENSBTAG00000033642 | N/A<br>N/A<br>N/A<br>N/A | 18<br>18<br>18<br>18 | 60703686-60969254<br>60459257-60466851<br>57640179-57681883<br>60781779-60803711 |
| Polydactyly | *ZNF423* | zinc finger protein 423 | Homo sapiens | ENSBTAG00000017397 | *ZNF423* | 18 | 18041277-18392207 |
| Polydactyly | *B9D1* | B9 domain containing 1 | Homo sapiens | ENSBTAG00000017790 | *B9D1* | 19 | 34709547-34717882 |

| Phenotype | Gene | Gene full name | Species | Gene ID | Bovine gene | BTA | Position |
|---|---|---|---|---|---|---|---|
| Syndactyly | *BHLHA9* | basic helix-loop-helix family member a9 | Homo sapiens | ENSBTAG00000048216 | *BHLHA9* | 19 | 22194038-22194712 |
| Syndactyly | *KCNJ2* | potassium voltage-gated channel subfamily J member 2 | Homo sapiens | ENSBTAG00000008294 | *KCNJ2* | 19 | 61185603-61195897 |
| Polydactyly | *MKS1* | Meckel syndrome, type 1 | Homo sapiens | ENSBTAG00000034963 | *MKS1* | 19 | 9378329-9393110 |
| Syndactyly | *SEPT9* | septin 9 | Homo sapiens | ENSBTAG00000002633 | *SEPT9* | 19 | 55118592-55153750 |
| Polydactyly | *Vps25* | vacuolar protein sorting 25 | Mus musculus | ENSBTAG00000019906 | *VPS25* | 19 | 43446967-43451724 |
| Polydactyly | *FBXW11* | F-box and WD repeat domain containing 11 | Homo sapiens | ENSBTAG00000015376 | *FBXW11* | 20 | 3584765-3624526 |
| Polydactyly | *MAP3K20* | mitogen-activated protein kinase kinase kinase 20 | Homo sapiens | ENSBTAG00000001083 | *ZAK* | 20 | 63111708-63127170 |
| Polydactyly | *KIF7* | kinesin family member 7 | Homo sapiens | ENSBTAG00000002440 | *KIF7* | 21 | 21482024-21496869 |
| Polydactyly | *MIPOL1* | mirror-image polydactyly 1 | Homo sapiens<br>Mus musculus<br>Rattus norvegicus<br>Bos taurus<br>Canis lupus familiaris<br>Capra hircus<br>Equus caballus<br>Felis catus<br>Ovis aries<br>Gallus gallus | ENSBTAG00000000655 | N/A | 21 | 47852815-48169316 |
| Syndactyly | *ADAMTS9* | a disintegrin-like and metallopeptidase (reprolysin type) with thrombospondin type 1 motif, 9 | Mus musculus | ENSBTAG00000003165 | *ADAMTS9* | 22 | 36876910-37037695 |
| Polydactyly | *IFT122* | intraflagellar transport 122 | Homo sapiens | ENSBTAG00000019121 | *IFT122* | 22 | 56881079-56948817 |
| Polydactyly | *LZTFL1* | leucine zipper transcription factor like 1 | Homo sapiens | ENSBTAG00000001484 | *LZTFL1* | 22 | 54104730-54115435 |
| Polydactyly | *ICK* | intestinal cell kinase | Homo sapiens | ENSBTAG00000015590 | *ICK* | 23 | 24982766-25027197 |

| Phenotype | Gene | Gene full name | Species | Gene ID | Bovine gene | BTA | Position |
|---|---|---|---|---|---|---|---|
| | *MBOAT1* | membrane bound O-acyltransferase domain containing 1 | Homo sapiens | ENSBTAG00000016519 | *MBOAT1* | 23 | 37498572-37608778 |
| Polydactyly | *Gata6* | GATA binding protein 6 | Mus musculus | ENSBTAG00000005734 | *GATA6* | 24 | 34549658-34579148 |
| Polydactyly | *IQCE* | IQ motif containing E | Homo sapiens | ENSBTAG00000014503 | *IQCE* | 25 | 41225570-41256752 |
| Polydactyly | *KIAA0556* | KIAA0556 | Homo sapiens | ENSBTAG00000006129 | *KIAA0556* | 25 | 25416302-25588540 |
| Syndactyly | *FBXW4* | F-box and WD repeat domain containing 4 | Homo sapiens | ENSBTAG00000003579 | *FBXW4* | 26 | 22228480-22311346 |
| Polydactyly, syndactyly | *FGFR2* | fibroblast growth factor receptor 2 | Homo sapiens | ENSBTAG00000014064 | *FGFR2* | 26 | 41823653-41926635 |
| Polydactyly | *Sufu* | SUFU negative regulator of hedgehog signaling | Mus musculus | ENSBTAG00000021068 | *SUFU* | 26 | 23452980-23517250 |
| Polydactyly | *TCTN3* | tectonic family member 3 | Homo sapiens | ENSBTAG00000011841 | *TCTN3* | 26 | 17000630-17021626 |
| Polydactyly, syndactyly | *FGFR1* | fibroblast growth factor receptor 1 | Homo sapiens | ENSBTAG00000015457 | *FGFR1* | 27 | 33250534-33291989 |
| Polydactyly | *BBS1* | Bardet-Biedl syndrome 1 | Homo sapiens | ENSBTAG00000020147 | N/A | 29 | 45200689-45215093 |
| Polydactyly, syndactyly | *DHCR7* | 7-dehydrocholesterol reductase | Homo sapiens | ENSBTAG00000016465 | *DHCR7* | 29 | 48930324-48950644 |
| Polysyndactyly | *FGF4* | fibroblast growth factor 4 | Mus musculus | ENSBTAG00000012563 | *FGF4* | 29 | 47656594-47657614 |
| Polydactyly | *TMEM138* | transmembrane protein 138 | Homo sapiens | ENSBTAG00000021604 | *TMEM138* | 29 | 40562910-40568784 |
| Polydactyly | *TMEM216* | transmembrane protein 216 | Homo sapiens | ENSBTAG00000021517 | *TMEM216* | 29 | 40586490-40590842 |
| Polydactyly | *CPLANE1* | ciliogenesis and planar polarity effector 1 | Homo sapiens | N/A | N/A | N/A | N/A |
| Polydactyly | *Dbf* | doublefoot | Mus musculus | N/A | N/A | N/A | N/A |
| Polydactyly | *DEL9P* | Chromosome 9p deletion syndrome | Homo sapiens | N/A | N/A | N/A | N/A |
| Syndactyly | *CUP2Q35* | Syndactyly, type I | Homo sapiens | N/A | N/A | N/A | N/A |
| Polydactyly | *DYNC2H1* | dynein cytoplasmic 2 heavy chain 1 | Homo sapiens | N/A | N/A | N/A | N/A |

| Phenotype | Gene | Gene full name | Species | Gene ID | Bovine gene | BTA | Position |
|---|---|---|---|---|---|---|---|
| Syndactyly | *EDSS2* | Ectodermal dysplasia-syndactyly syndrome 2 | Homo sapiens | N/A | N/A | N/A | N/A |
| Polydactyly | *Fpdty* | forelimb polydactyly | Mus musculus | N/A | N/A | N/A | N/A |
| Polydactyly | *Hpdty* | hindlimb polydactyly | Mus musculus | N/A | N/A | N/A | N/A |
| Polysyndactyly | *HOXA@* | homeobox A cluster | Homo sapiens | N/A | N/A | N/A | N/A |
| Polydactyly | *LOC106049962* | SHH silencer LOC106049962 | Homo sapiens | N/A | N/A | N/A | N/A |
| Syndactyly | *MSSD* | syndactyly, mesoaxial synostotic, with phalangeal reduction | Homo sapiens | N/A | N/A | N/A | N/A |
| Polydactyly | *PAPA2* | postaxial polydactyly, type A2 | Homo sapiens | N/A | N/A | N/A | N/A |
| Polydactyly | *PAPA3* | Polydactyly, postaxial, type A3 | Homo sapiens | N/A | N/A | N/A | N/A |
| Polydactyly | *PAPA4* | Polydactyly, postaxial, type A4 | Homo sapiens | N/A | N/A | N/A | N/A |
| Polydactyly | *PAPA5* | Polydactyly, postaxial, type A5 | Homo sapiens | N/A | N/A | N/A | N/A |
| Polydactyly | *pcp* | polydactyly with cleft palate | Mus musculus | N/A | N/A | N/A | N/A |
| Polydactyly | *Plsm1* | Polydactyly-luxate syndrome (PLS) morphotypes QTL 1 | Rattus norvegicus | N/A | N/A | N/A | N/A |
| Polydactyly | *Plsm2* | Polydactyly-luxate syndrome (PLS) morphotypes QTL 2 | Rattus norvegicus | N/A | N/A | N/A | N/A |
| Polydactyly | *Plsm3* | Polydactyly-luxate syndrome (PLS) morphotypes QTL 3 | Rattus norvegicus | N/A | N/A | N/A | N/A |
| Polydactyly | *Plsm4* | Polydactyly-luxate syndrome (PLS) morphotypes QTL 4 | Rattus norvegicus | N/A | N/A | N/A | N/A |
| Polydactyly | *Po* | postaxial polydactyly | Mus musculus | N/A | N/A | N/A | N/A |
| Polysyndactyly | *PS* | polysyndactyly | Mus musculus | N/A | N/A | N/A | N/A |
| Polydactyly | *py* | polydactyly | Mus musculus | N/A | N/A | N/A | N/A |
| Polydactyly | *T1pdty* | type I polydactyly | Mus musculus | N/A | N/A | N/A | N/A |
| Polydactyly | *TBX1* | T-box 1 | Homo sapiens | N/A | N/A | N/A | N/A |
| Polydactyly | *TMEM231* | transmembrane protein 231 | Homo sapiens | N/A | N/A | N/A | N/A |
| Polydactyly | *WDR35* | WD repeat domain 35 | Homo sapiens | N/A | N/A | N/A | N/A |
| Polydactyly | *Xpl* | X-linked polydactyly | Mus musculus | N/A | N/A | N/A | N/A |

| Phenotype | Gene | Gene full name | Species | Gene ID | Bovine gene | BTA | Position |
|---|---|---|---|---|---|---|---|
| Polydactyly, polysyndactyly | *ZRS* | ZPA regulatory sequence | Homo sapiens<br>Mus musculus | N/A | N/A | N/A | N/A |
| Polydactyly | *Arhgap6* | Rho GTPase activating protein 6 | Mus musculus | ENSBTAG00000002626 | *ARHGAP6* | X | 137588934-137693777 |
| Syndactyly | *BCOR* | BCL6 corepressor | Homo sapiens | ENSBTAG00000047339 | *BCOR* | X | 108883915-108907652 |
| Syndactyly | *CCNQ* | cyclin Q | Homo sapiens | ENSBTAG00000011350 | *CCNQ* | X | 39727312-39736823 |
| Polydactyly | *GPC3* | glypican 3 | Homo sapiens | ENSBTAG00000020406 | *GPC3* | X | 17305874-17770661 |
| Polydactyly | *GPC4* | glypican 4 | Homo sapiens | ENSBTAG00000020644 | *GPC4* | X | 17087447-17122426 |
| Polydactyly | *Hccs* | holocytochrome c synthetase | Mus musculus | ENSBTAG00000012113 | *HCCS* | X | 137706223-137714932 |
| Polydactyly | *Mid1* | midline 1 | Mus musculus | ENSBTAG00000010152 | *MID1* | X | 139791005-139928580 |
| Polydactyly | *Msl3* | MSL complex subunit 3 | Mus musculus | ENSBTAG00000009278 | *MSL3* | X | 142041442-142056938 |
| Syndactyly | *NAA10* | N(alpha)-acetyltransferase 10, NatA catalytic subunit | Homo sapiens | ENSBTAG00000047702 | *NAA10* | X | 40058809-40063513 |
| Polydactyly, syndactyly | *OFD1* | OFD1, centriole and centriolar satellite protein | Homo sapiens | ENSBTAG00000004461<br>ENSBTAG00000048102 | *OFD1*<br>N/A | X<br>X | 136925770-136987149<br>143736738-143800954 |
| Syndactyly | *PORCN* | porcupine O-acyltransferase | Homo sapiens | ENSBTAG00000009282 | *PORCN* | X | 91723575-91734163 |

**Table S2.** Primer pairs used for validation of the two variants (g.12552417dupC, g.12552582delA) in *XIRP1* by Sanger sequencing of the 898 bp amplicon and genotyping by RFLP. Primer pairs sequences, length, annealing temperature (AT), amplicon size (AS) in base pairs (bp), restriction enzyme, buffer and incubation temperature (IT) are given.

| **Primer (F=forward, R=reverse, mism=mismatch)** | **Target** | **Variant** | **Primer sequence (5′ - 3′)** | **Length** | **AT (°C)** | **AS (bp)** | **Restrictions enzyme** | **Buffer** | **IT (°C)** |
|---|---|---|---|---|---|---|---|---|---|
| BTA_XIRP1_F1 | gDNA | g.12552582delA | GCTGCATAAACAAAGGAGACC | 21 | 60 | | | | |
| BTA_XIRP1_R1 | | | AGGCGTTACGAGAGTGCAGA | 20 | 60 | 898 | *Xma*I | Cut-Smart | 37 |
| BTA_XIRP1_Fmism | gDNA | g.12552417dupC | AAAGCACTGCCTGGAGGGCTAT CTGGCAGG | 30 | 62 | | | | |
| BTA_XIRP1_R1 | | | AGGCGTTACGAGAGTGCAGA | 20 | 62 | 341 | *Bsp*MI | NEBuffer 3.1 | 37 |

**Table S3.** Primer sequences used for NGS data validation of the ZRS and pZRS region of *LMBR1* gene using sanger sequencing. Primer pairs, length, annealing temperature (AT) and amplicon size (AS) in base pairs (bp) and are given.

| Primer (Intron) (F=forward, R=reverse) | Target | Primer sequence (5′ - 3′) | Length | AT (°C) | AS (bp) |
|---|---|---|---|---|---|
| LMBR1_Int5_F1 | gDNA | ACTGCACAGACTCTGAACATG | 21 | 58 | |
| LMBR1_Int5_R1 | | ACGTAAGATGCAAGCAGAGG | 20 | 58 | 930 |
| LMBR1_Int5_F2 | gDNA | GTGGTTTTGATAAAGCAGGCT | 21 | 58 | |
| LMBR1_Int5_R2 | | CATTTGGGGATGTCTGGATG | 20 | 58 | 853 |
| LMBR1_Int5_F3 | gDNA | GATGACAGATTATTTCATGAACC | 23 | 58 | |
| LMBR1_Int5_R3 | | AGCCCTGTACGTCACAACAC | 20 | 58 | 992 |

**Table S4.** Results of the filtered whole genome sequencing data. All variants, which are homozygous or heterozygous mutant exclusively in the affected polydactylous calves combined with heterozygous mutant or homozygous wild type allels in the dam of Case 2. The remaining critical variants for the phenotype polydactyly and polysyndactyly are printed in bold.

| Gene | BTA | Position | ID | Base change | Genotype | | | cDNA | Protein | Transcript | PolyPhen |
|---|---|---|---|---|---|---|---|---|---|---|---|
| | | | | | Case 1 | Case 2 | Dam of Case 2 | | | | |
| *C7orf72* | 4 | 5672513 | - | A>AG | 0/1 | 0/1 | 0/1 | c.1088_1089insC | p.Arg364fs | ENSBTAT00000061416.2 | Benign 0.106 |
| *CERK* | 5 | 118035410 | - | CACTCGGTGCACTGCACGGAGAACACACGCGTGAGCCCCGCACCACGCCAGCTCCCCTGACCGCCCCTT>C | 0/1 | 0/1 | 0/0 | c.119-58_128delAAGGGGCGGTCAGGGGAGCTGGCGTGGTGCGGGGCTCACGCGTGTGTTCTCCGTGCAGTGCACCGAGT | p.Val40fs | ENSBTAT00000066162.1 | Probably damaging 1.000 |
| *ENSBTAG00000047383* | 12 | 71848598 | - | GCACC>G | 0/1 | 0/1 | 0/0 | c.3082_3085delGGTG | p.Gly1028fs | ENSBTAT00000050045.3 | Possibly damaging 0.891 |
| *ENSBTAG00000047383* | 12 | 71848603 | - | A>AATGG | 0/1 | 0/1 | 0/0 | c.3080_3081insCCAT | p.Gly1028fs | ENSBTAT00000050045.3 | Possibly damaging 0.536 |
| ***XIRP1*** | **22** | **12552417** | **-** | **A>AC** | **0/1** | **0/1** | **0/1** | **c.3387dupG** | **p.Trp1130fs** | **ENSBTAT00000065632.1** | **Possibly damaging 0.922** |
| ***XIRP1*** | **22** | **12552582** | **-** | **CA>C** | **0/1** | **0/1** | **0/0** | **c.3222delT** | **p.Ala1076fs** | **ENSBTAT00000065632.1** | **Probably damaging 0.998** |

**Table S5.** Genotyping results of the filtered variants (g.12552417dupC, g.12552582delA) in *XIRP1* in 359 cattle of 8 different breeds using restriction fragment length polymorphisms (RFLP). The distribution of the wild type allele (wt) and mutant allele (mut) is shown. Eight animals of the Holstein breed were heterozygous and one homozygous carrier of the compound of the variants g.12552417dupC and g.12552582delA. None of the variants were detected in the other breeds.

| | | **g.12552417dupC (*XIRP1*)** | | | **g.12552582delA (*XIRP1*)** | | |
|---|---|---|---|---|---|---|---|
| **Breeds** | **Number of animals** | **Genotype wt/wt** | **Genotype wt/mut** | **Genotype mut/mut** | **Genotype wt/wt** | **Genotype wt/mut** | **Genotype mut/mut** |
| Holstein | 276 | 267 | 8 | 1 | 267 | 8 | 1 |
| Fleckvieh | 12 | 12 | 0 | 0 | 12 | 0 | 0 |
| Brown | 12 | 12 | 0 | 0 | 12 | 0 | 0 |
| Salers | 12 | 12 | 0 | 0 | 12 | 0 | 0 |
| German Angus | 12 | 12 | 0 | 0 | 12 | 0 | 0 |
| Charolais | 12 | 12 | 0 | 0 | 12 | 0 | 0 |
| Limousin | 11 | 11 | 0 | 0 | 11 | 0 | 0 |
| Blonde d’Aquitaine | 12 | 12 | 0 | 0 | 12 | 0 | 0 |
