## Supplementary Figures 1-4 for "*De-novo* variants in *XIRP1* associated with polydactyly and polysyndactyly in Holstein cattle"

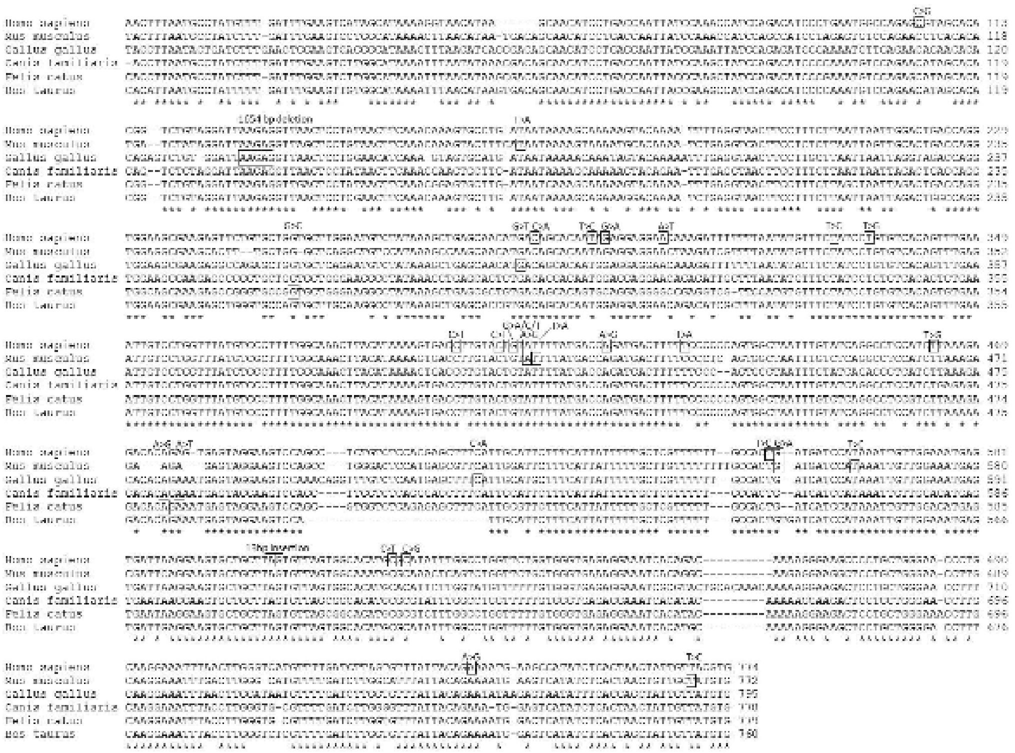

**Figure S1.** Sequence of the bovine ZRS region in *LMBR1* gene matched to Homo sapiens, Mus musculus, Gallus gallus, Canis familiaris and Felis catus. Previously reported variants, which are associated with polydactyly in the different species, are framed. Identical sequences between the species are marked with an asterisk.

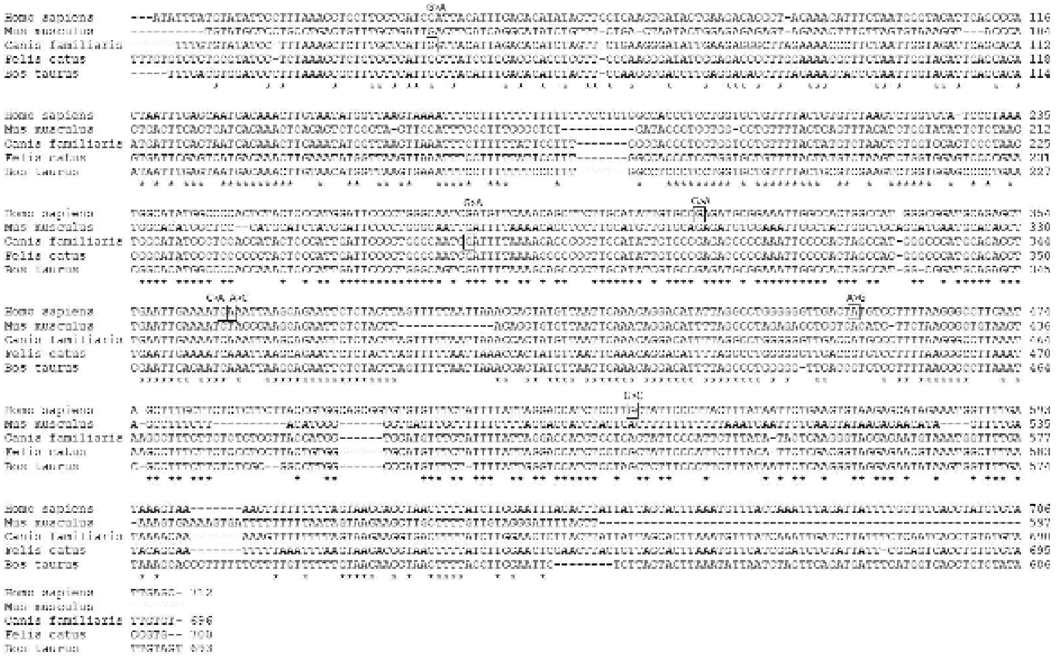

**Figure S2.** Sequence of the bovine pZRS region in *LMBR1* gene matched to Homo sapiens, Mus musculus, Canis familiaris and Felis catus. Previously reported variants, which are associated with polydactyly in the different species, are framed. Identical sequences between the species are marked with an asterisk.

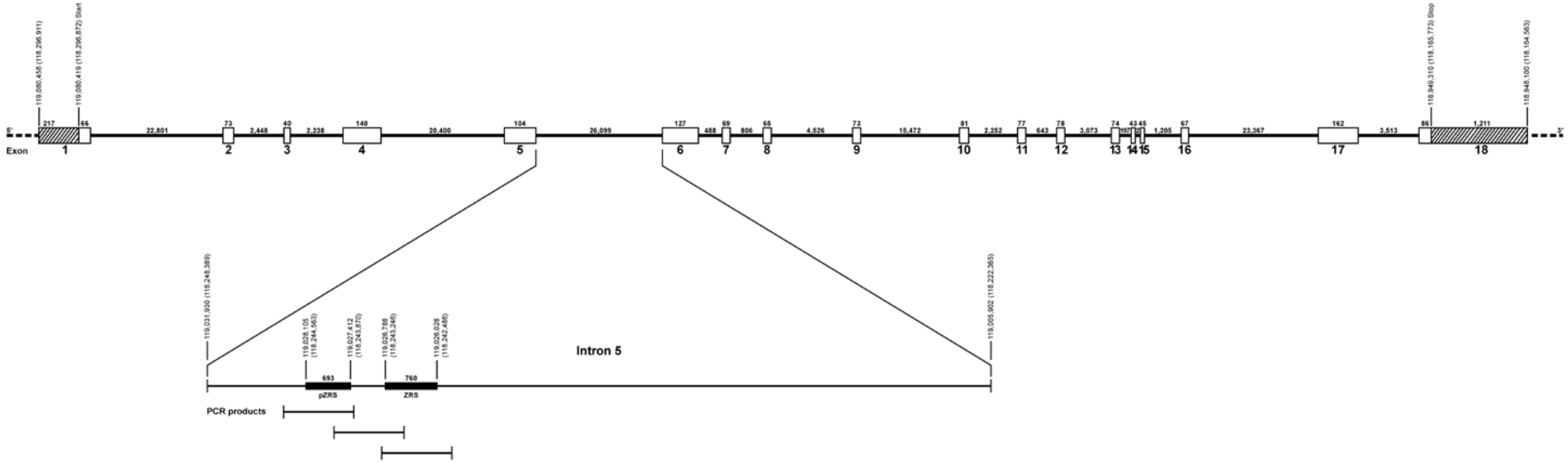


**Figure S3.** Gene model of the bovine *LMBR1* gene (ENSBTAG00000030817, Transcript ENSBTAT00000043570.3/ ENSBTAT00000043570.4) located on chromosome 4 according to UMD3.1 and ARS-UCD1.2. The pZRS and ZRS region in intron 5 is marked in bold. For variant analysis by sanger sequencing, three overlapping amplicons were produced. The first location information were according to UMD3.1 and the second in parentheses were accordin to ARS-UCD1.2.

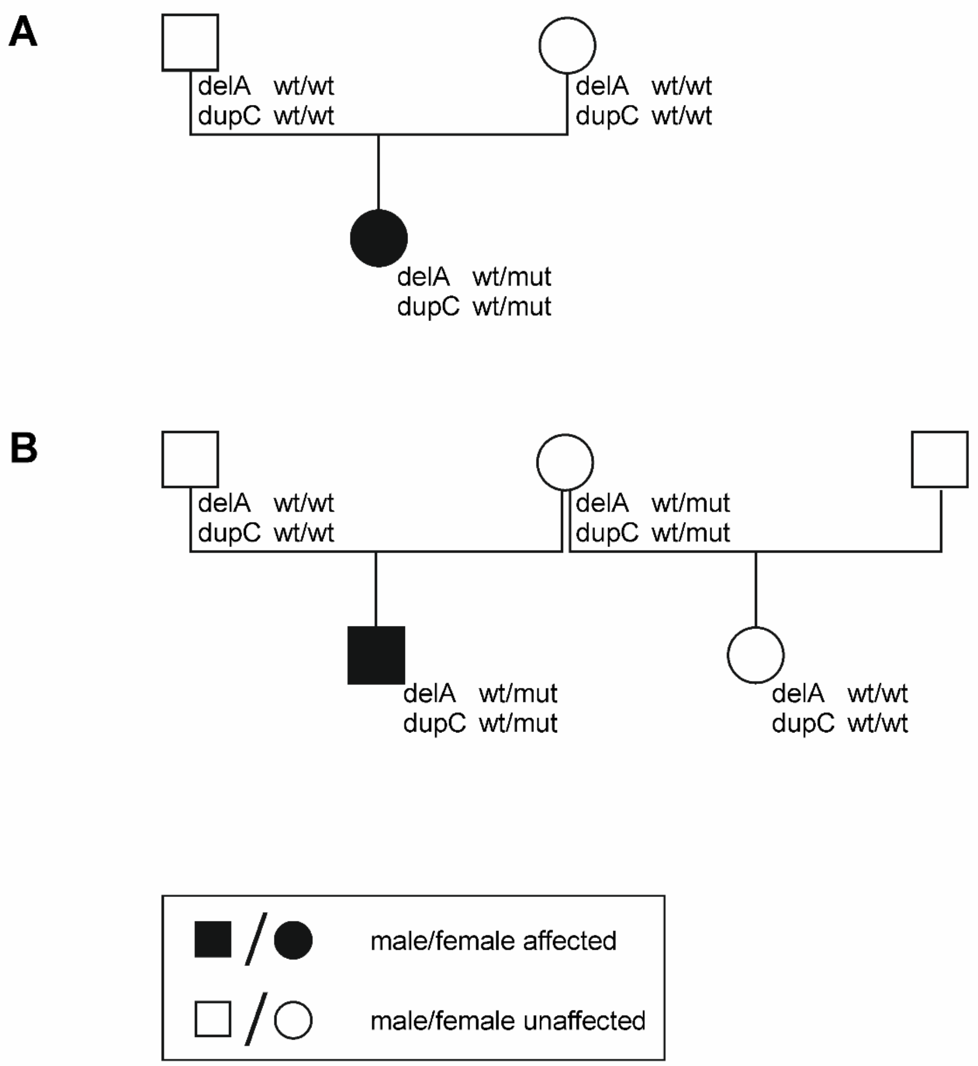
A
delA wt/wt
dupC wt/wt
delA wt/wt
dupC wt/wt
delA wt/mut
dupC wt/mut
B
delA wt/wt
dupC wt/wt
delA wt/mut
dupC wt/mut
delA wt/mut
dupC wt/mut
delA wt/wt
dupC wt/wt
male/female affected
male/female unaffected

**Figure S4.** Validation results of the familial animals of the two polydactylous calves. (A) edigree of the first female affected calf (case 1) and (B) pedigree of the second male affected calf (case 2). The wild type (wt) and mutant (mut) occurrence of the variants (g.12552417dupC, g.12552582delA) were added to the pedigree for each genotyped animal, were samples were available.
